# Individual differences shape conceptual representation in the brain

**DOI:** 10.1101/2025.08.22.671848

**Authors:** Matteo Visconti di Oleggio Castello, Tom Dupré la Tour, Jack L. Gallant

## Abstract

Each person experiences the world through a unique conceptual lens, shaped by personal experiences, natural variations, or disease. These individual differences have remained largely inaccessible to cognitive neuroscience and clinical neurology, limiting the development of precision medicine approaches to cognitive disorders. To overcome this limitation, here we develop a new statistical framework to measure and interpret individual differences in functional brain representations. We apply this framework to characterize how different individuals represent the same concepts. Twenty-four participants listened to narrative stories while their brain activity was measured with functional MRI (fMRI). Encoding models were used to recover how hundreds of concepts were represented in each person’s brain. Despite listening to identical stories, participants showed systematic individual differences in conceptual representations. These differences reveal person-specific biases in how concepts are represented in the brain. Individual variability was highest in regions that represent social information. Because these regions are thought to integrate sensory information with personal beliefs and experiences, the observed individual differences may reflect cognitive traits unique to each person. Our work reveals that individual differences are a systematic, measurable principle of conceptual representations in the human brain. By enabling researchers to measure and interpret differences in person-specific functional brain representations, our work establishes a new paradigm for precision neuroscience. This paradigm provides a rigorous foundation for developing fMRI applications in precision medicine to diagnose and monitor cognitive disorders.

## Introduction

Brain function varies across individuals due to differences in brain development and age, or due to disease (Insel & Cuthbert, 2015; Salthouse, 2019). Individual differences manifest naturally in many cognitive functions, including attention (Hunt et al., 1989; Wager et al., 2005), working memory (Jarrold & Towse, 2006; Rypma & D’Esposito, 1999), decision-making (Engelmann & Tamir, 2009; Newman et al., 2003), language processing (Kidd et al., 2018; King & Just, 1991), and semantic memory (Chadwick et al., 2016; Hoffman, 2018). The same cognitive functions are critically impacted by several cognitive syndromes and disorders (Bang et al., 2015; Barch, 2005; Frith & Happé, 2005; Perry et al., 2000). To improve the diagnosis and monitoring of clinical disorders, it is necessary to distinguish non-pathological from pathological variations in brain function. However, individual differences are rarely studied in cognitive neuroscience, in neuroimaging, or in clinical neurology. Instead, these fields commonly focus on group-level effects. This approach treats individual variations as a nuisance rather than as a source of meaningful information, and thus impedes the translation of neuroimaging findings to individualized clinical diagnoses. Shifting the focus from group-level effects to individual differences would clarify how individual variation in brain function impacts behavior, and would provide a platform for making individualized clinical diagnoses based on neuroimaging data (Insel & Cuthbert, 2015; Woo et al., 2017).

One important roadblock to examining individual differences in brain function is the lack of a systematic and principled statistical framework for detecting, measuring, and interpreting such differences. Neuroimaging studies are commonly based on statistical methods and experimental paradigms that are optimized to detect group-level effects, discarding information necessary for detecting individual differences. In a typical study, participants perform an experimental task while their brain activity is measured with functional MRI (fMRI). To draw general conclusions about the relationship between the task and brain activity, the results are usually analyzed at the group level. Each participant’s data are spatially transformed to a common anatomical brain space, and all modeling is performed on this spatially transformed data. The modeling results are then averaged across participants to increase statistical power and draw conclusions based on the group of participants. Spatially transforming and averaging the data across participants discards any individual variations that exist either in brain anatomy or in brain activity.

The limitations of group-level studies are well known (Fisher et al., 2018; Ince et al., 2021, 2022; McManus et al., 2023), and some neuroscience researchers have argued in favor of *precision* studies (Huth et al., 2012, 2016; Lee et al., 2024; Michon et al., 2022; Naselaris et al., 2021; Poldrack, 2017) in which a large amount of data is collected in each individual to support participant-specific analyses. One common approach to precision neuroimaging is to measure an individual’s brain activity while they perform a highly controlled task under a few different task conditions (Barch et al., 2013; Botch & Finn, 2024; Chadwick et al., 2016; Charest et al., 2014; Engelmann & Tamir, 2009; Grabner et al., 2007; Newman et al., 2003; Rypma & D’Esposito, 1999). However, such narrow tasks can only reveal individual differences along the small number of conditions that were manipulated in the experiment. In many cases these targeted dimensions will neither be those that show the largest individual differences, nor those that provide the best diagnostic information. Thus, conventional task-based experiments are both insufficient and inefficient for reliably detecting individual differences in cognitive function.

Another common approach to precision neuroimaging focuses on resting-state functional connectivity (Biswal et al., 1995; Greicius et al., 2003; Yeo et al., 2011). In these experiments, brain activity is measured while participants lie in the scanner without being given any explicit task or instruction. The measured fMRI data are used to identify networks of brain areas that exhibit temporally correlated activity. Precision resting state functional connectivity studies have examined individual variations in the strength of these temporal correlations or in the spatial organization of the putative networks (Feilong et al., 2018, 2021; Finn et al., 2015, 2020; Finn & Bandettini, 2021; Gordon et al., 2015, 2017; Gratton et al., 2018; Mueller et al., 2013; Vanderwal et al., 2017). However, in the absence of an explicit task it is impossible to identify and characterize the factors that cause individual variability in functional connectivity networks (Meschke et al., 2023), and it is impossible to examine how representations of specific information differ across individuals. Thus, resting-state functional connectivity experiments are insufficient for explaining individual differences in cognitive function and representations.

The limitations of conventional group-based and precision neuroimaging studies highlight a pressing need for a new experimental and analytical framework for studying individual differences in brain function. From a theoretical standpoint, such a framework must use an experimental paradigm that can reliably detect individual differences in complex functional representations, and it must implement a formal statistical approach for measuring and interpreting these individual differences.

Here we present a new framework that addresses both of these requirements, and we use this framework to measure and interpret individual differences in the representation of conceptual information conveyed by natural speech. The lexical-semantic representations that we characterize in this study are found within multimodal brain regions (Binder et al., 2009; Binder & Desai, 2011; Damasio & Damasio, 1994; Deniz et al., 2019; Huth et al., 2016; Martin & Chao, 2001; Patterson et al., 2007) that are part of a distributed conceptual network of unimodal, multimodal, and executive areas (Çukur et al., 2013; Gallant & Popham, 2020; Huth et al., 2012; LeBel et al., 2021). This distributed conceptual network is severely impacted by many cognitive and neurological disorders, including Alzheimer’s disease (Parasuraman & Haxby, 1993; Perry et al., 2000), frontotemporal dementia (Bang et al., 2015; Ranasinghe et al., 2016), schizophrenia (Barch, 2005; Mitchell et al., 2001), and autism spectrum disorder (Dichter, 2012; Frith & Happé, 2005). In this study we focus specifically on how lexical-semantic representations differ across healthy young adults. This information will necessarily underpin any future studies of cognitive syndromes and neurological disorders that affect conceptual processing.

## Results

### Recovering cortical representations of conceptual information in individual people

Individual differences in conceptual representation remain largely unexplored. This gap exists partly because conventional neuroimaging studies typically use a small number of experimental conditions, and so each study can only reveal individual differences across a narrow set of concepts. To efficiently examine individual differences across many conceptual representations, we used a naturalistic experimental paradigm that evokes hundreds of distinct lexical-semantic representations. Twenty-four participants listened to eleven autobiographical stories from *The Moth Radio Hour* podcast while their brain activity was measured with fMRI (Huth et al., 2016). Each story was narrated by a different storyteller who described personal life events, conveying complex information associated with many lexical-semantic concepts.

The Voxelwise Encoding Model (VEM) framework (Dupré la Tour et al., 2025; Naselaris et al., 2011; Wu et al., 2006) was used to model brain activity evoked by the narrative stories, separately in each individual participant (Figure 1A). In the VEM framework, one or more feature spaces are first extracted from the experimental stimulus. These feature spaces quantify some specific type of information that is hypothesized to be represented in brain activity, such as lexical-semantics or low-level properties of speech. Regularized linear regression is then used to create voxelwise encoding models that predict brain activity from these feature spaces. If a voxelwise encoding model predicts brain activity significantly, it can be concluded that information contained within the feature spaces is also represented in the brain. Finally, the encoding model weights are plotted on the cortical surface to map the spatial organization of the specific features that are represented in the brain.

**Figure 1.**
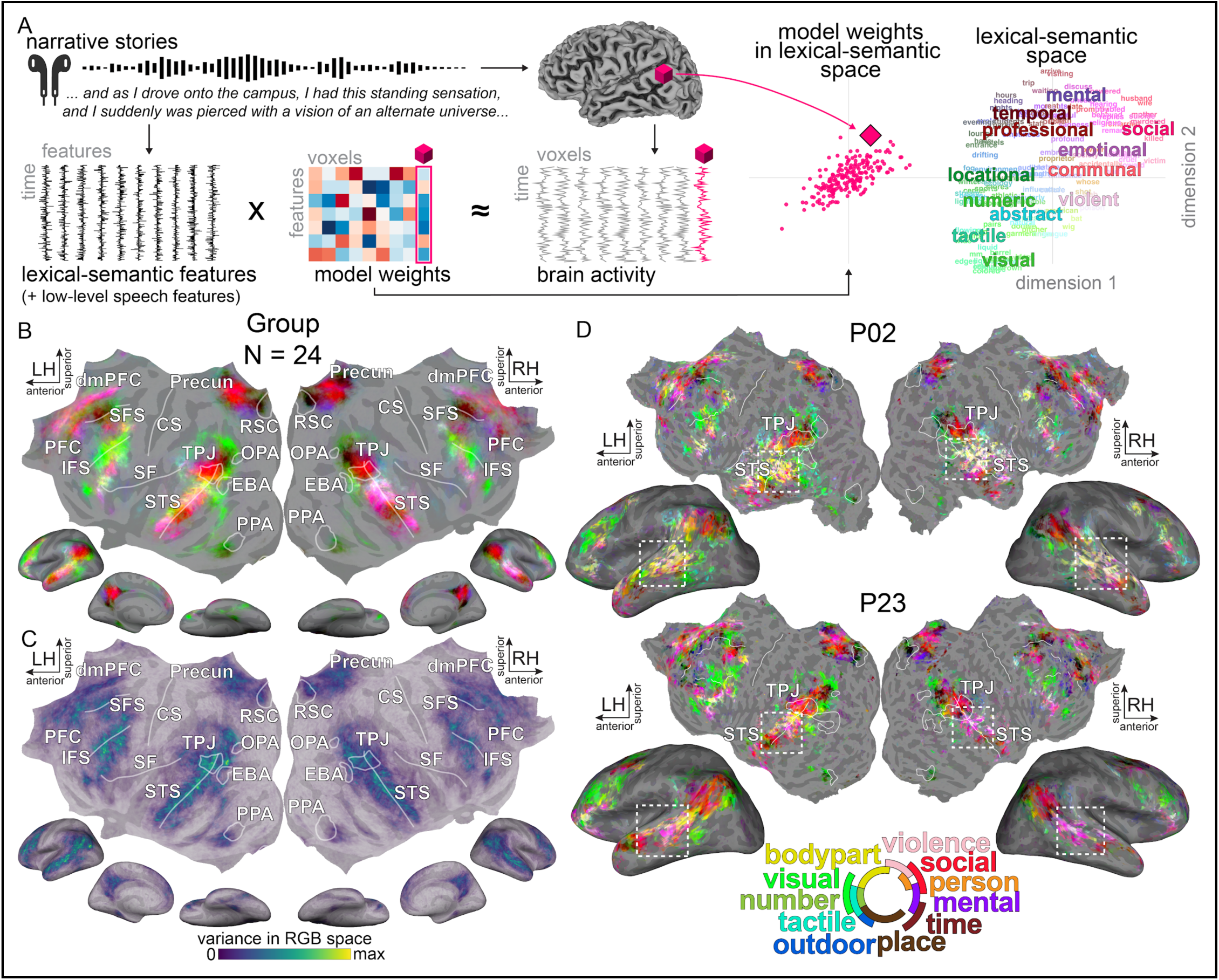
Recovering individual-specific cortical representations of lexical-semantic concepts. To investigate individual differences in conceptual representation, we used a naturalistic paradigm that evokes a large number of distinct lexical-semantic representations. [A] Twenty-four participants listened to over two hours of narrative stories while their brain activity was measured with functional MRI (fMRI). For each participant, a separate voxelwise encoding model was used to predict fMRI responses from high-dimensional lexical-semantic features extracted from the stories. The model weights for each voxel can be considered geometrically as a single point in the high-dimensional lexical-semantic space. The location of that point determines the lexical-semantic concept represented in the brain activity measured from that voxel. [B] Cortical maps of lexical-semantic representations are created by projecting high-dimensional model weights onto a reduced 3-dimensional lexical-semantic space (see Methods). A continuous RGB colormap is used to color each voxel according to the projection onto the 3D lexical-semantic space, such that different colors indicate representations of different lexical-semantic concepts. The cortical map shows the group-averaged lexical-semantic map (N=24). This group-averaged map reveals cortical areas that represent lexical-semantic concepts consistently across all participants, but necessarily obscures potential variability across individuals. [C] To highlight this variability, the cortical map shows the variance of the participant-specific lexical-semantic maps. Regions around the temporo-parietal junction (TPJ), superior temporal sulcus (STS) and portions of the frontal cortex show the highest variance across participants. These high-variance regions suggest systematic individual differences that warrant further investigation. [D] Individual-specific lexical-semantic maps are shown for two representative participants (see Supplementary Figure 3 for all participants). Dashed squares highlight regions with variable lexical-semantic representations across participants. These participant-specific maps demonstrate individual differences in lexical-semantic representations.

In the study described here, we wish to determine how lexical-semantic information is represented in the brain of each individual participant. Therefore, a lexical-semantic feature space was created by projecting each word in the stories onto a 985-dimensional word embedding space (*english1000*; Deniz et al., 2019; Huth et al., 2016; LeBel et al., 2021, 2023). An additional low-level speech feature space was included to account for variability in the speech pattern of different speakers. These two feature spaces were used jointly to estimate a single voxelwise encoding model for each participant. This model was fit on a training set consisting of fMRI data measured in response to ten different stories, and it was evaluated on a separate held-out test set consisting of a new story repeated multiple times (see Methods). Inspection of the model prediction accuracy revealed that the lexical-semantic encoding models predict brain activity significantly better than chance within temporal, parietal, and frontal regions (see Supplementary Figure 1). This finding confirms that these voxelwise encoding models reliably capture lexical-semantic representations across a distributed network of brain areas involved in the representation of multimodal conceptual knowledge (Binder et al., 2009; Binder & Desai, 2011; Damasio & Damasio, 1994; Deniz et al., 2019; Gallant & Popham, 2020; Huth et al., 2016; Martin & Chao, 2001; Patterson et al., 2007).

To identify how these lexical-semantic representations are spatially organized, and to determine whether they are variable across individuals, we sought to examine the pattern of encoding model weights across the cortical surface. Because the lexical-semantic model weights are 985-dimensional, it is difficult to visualize all dimensions on the cortical surface. Therefore, for visualization purposes the model weights in each voxel were first projected onto a reduced three-dimensional lexical-semantic space (Huth et al., 2016; see Methods). Then, the three dimensions were mapped onto the RGB color space to plot the selectivity of each voxel in this reduced lexical-semantic space. This visualization procedure produces a continuous colormap in which different colors indicate different lexical-semantic representations. For example, red indicates representations of social concepts, and green indicates representations of visual concepts (Figure 1).

Figure 1B shows the group-averaged lexical-semantic map across the 24 participants. This map reveals several cortical regions that represent lexical-semantic concepts in a way that is consistent with findings from prior studies. For example, the temporo-parietal junction (TPJ) and precuneus (Precun) underpin social cognition and theory-of-mind processes (Gobbini et al., 2007; Saxe & Kanwisher, 2003; Tamir et al., 2016). Accordingly, these areas are colored in red in the lexical-semantic map, indicating representations of social concepts.

Similarly, the ventral-temporal cortex is involved in the representation of visual object categories (Haxby et al., 2001; Margalit et al., 2020). Accordingly, patches in the ventral-temporal cortex are colored in green, indicating representations of visual concepts. Although the group-averaged map reveals a broad consistency of lexical-semantic selectivity across participants, it obscures potential individual variability. To highlight this variability, Figure 1C shows the variance of the lexical-semantic maps across participants. This variance map reveals that regions near the temporo-parietal junction (TPJ), superior temporal sulcus (STS), and portions of the frontal cortex show the highest variance across participants. This high variance suggests that these regions may exhibit substantial individual variability in lexical-semantic representations. To observe this variability more directly, Figure 1D shows the individual-specific lexical-semantic maps of two representative participants (see Supplementary Figure 3 for all participants). Consistent with the variance map, inspection of the individual-specific lexical-semantic maps reveals considerable individual differences within the bilateral STS. The STS of P02 is colored in green and orange, indicating representations of visual and people-related concepts. In contrast, the STS of P23 is colored in red and purple, indicating representations of social and mental-related concepts. The variability in the lexical-semantic maps suggest that individuals differ in how they represent conceptual information when listening to the same narrative stories.

### A novel statistical framework for modeling individual differences in functional representations

The cortical maps in Figure 1 reveal the existence of individual differences in lexical-semantic representations. However, these maps are based on a reduced three-dimensional lexical-semantic space for visualization purposes. This reduced space explains on average only 21.45% of the total variance in each participant’s model weights (95% confidence interval [CI]: 20.25% to 22.58%, bootstrap CI). Because these cortical maps necessarily discard almost 80% of the variance in the model weights of each participant, individual differences in lexical-semantic representations are likely larger than those suggested by inspecting the cortical maps.

To examine individual differences in the full lexical-semantic space, in this study we developed a novel statistical approach based on the mathematical theory of optimal transport (Peyré & Cuturi, 2019). In this approach, the encoding model weights in each voxel are considered geometrically as a point in the 985-dimensional lexical-semantic space (Figure 2A). For each participant, the collection of all lexical-semantic weights is modeled as a 985-dimensional Gaussian distribution in the lexical-semantic space. Thus, each participant is associated with their own distribution of lexical-semantic model weights. Individual differences in lexical-semantic representations are formalized as differences in these high-dimensional Gaussian distributions (see Methods). Each participant’s distribution of lexical-semantic model weights consists of a unique participant-specific component and a group component that is shared across participants. To examine individual variability across participants, the individual and group components must first be disentangled. To do this, each participant’s distribution of model weights is transformed to an independent group distribution that includes the model weights of all other participants (leave-one-participant-out procedure, Figure 2B). This transformation is achieved by means of the optimal transport (Figure 2C). Optimal transport ensures that each participant’s model weights are transformed minimally in the lexical-semantic space, while simultaneously maximizing the similarity between the covariances of the participant’s transformed distribution and the group distribution (see Methods). Once the participant’s model weights have been transformed, individual differences between the participant-specific weights and the group distribution can be identified. These differences can be visualized in a low-dimensional space for interpretation purposes. Alternatively, the magnitude of the total difference in model weights between an individual participant and the group can be expressed as a simple summary statistic, the distributional explained variance (dEV). The dEV is a generalization of the measure of variance explained by a subspace (see Methods), and it can be interpreted as the fraction of variance in the participant’s model weights that is explained by the group distribution.

**Figure 2.**
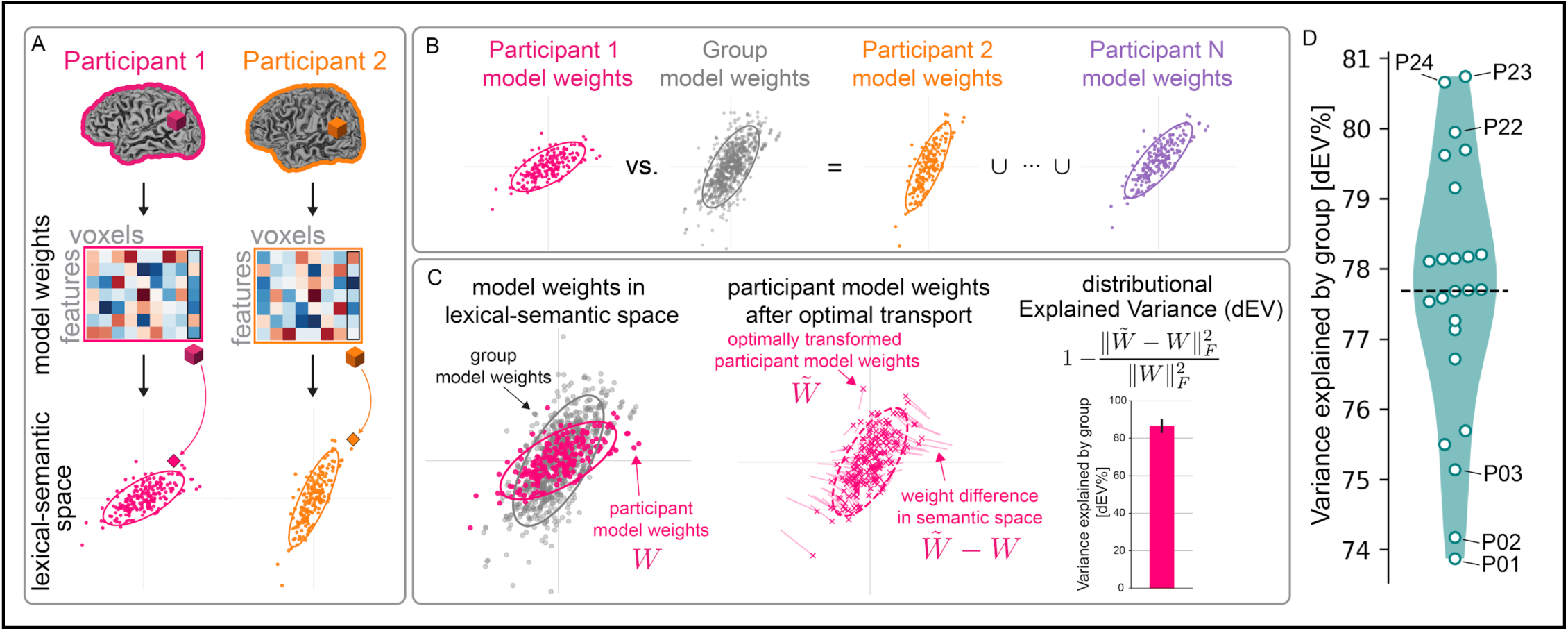
A novel statistical framework for modeling individual differences in functional representations. We developed a framework based on optimal transport theory to measure and interpret individual differences in representations recovered by voxelwise encoding models. [A] For each participant the collection of all voxelwise model weights is modeled as a high-dimensional Gaussian distribution in the lexical-semantic space. This distribution reflects the lexical-semantic concepts represented by each participant’s brain activity during the experiment. [B] To determine individual differences in these distributions of model weights, each participant’s distribution is compared to an independent group distribution that includes the model weights of all other participants. [C] Optimal transport is used to transform each participant’s model weights to maximize the similarity with the group distribution, while minimizing changes in lexical-semantic space. The distributional explained variance (dEV) is then computed from these optimally transformed weights to quantify the fraction of variance in a participant’s model weights that is explained by the group distribution. [D] Distribution of dEV values across the 24 participants in the story-listening experiment. The dashed line shows the group average (77.68% dEV). The group distribution explains on average ∼78% of the variance in the model weights of individual participants, indicating that the lexical-semantic representations are largely consistent during narrative story-listening. However, ∼22% of variance cannot be explained by the group distribution. This unexplained variance reflects individual differences in lexical-semantic representations.

In this study, the dEV was used to measure the extent of individual differences in lexical-semantic representations during the narrative story-listening experiment (Figure 2D). The average dEV across participants is 77.68% (95% CI: 76.92% to 78.39%), meaning that on average 78% of the variance in a participant’s lexical-semantic model weights is explained by the group model. Thus, lexical-semantic representations are largely consistent across individuals. However, over 20% of the variance in each participant’s lexical-semantic model weights cannot be explained by the group distribution. This unexplained variance reflects individual differences in lexical-semantic representations.

### Spatial heterogeneity of individual differences in lexical-semantic representations

Prior work has suggested that individual differences in functional organization are spatially heterogeneous, with greater individual variability reported in association and cognitive areas than in sensory areas (Cui et al., 2020; Frost & Goebel, 2012; Gratton et al., 2018; Mueller et al., 2013; Vanderwal et al., 2017; Zilles & Amunts, 2013). However, these findings were derived from resting-state data or simple controlled experiments that did not assess natural language comprehension. Therefore, it remains unknown whether individual variability of complex lexical-semantic representations exhibits similar spatial heterogeneity across the cortex.

Because lexical-semantic representations are distributed broadly across the cortex (Figure 1), we hypothesized that lexical-semantic representations in higher-order cognitive regions exhibit greater individual variability than representations in regions close to unimodal sensory systems. Testing this hypothesis requires a measure of individual variability for each separate voxel. To generate this measure, we first modified the dEV summary statistic to obtain a dEV value for each voxel (see Methods). Because the dEV quantifies the amount of variance that is explained by the group, we then expressed individual variability as 1 - dEV, which corresponds to the amount of variance unexplained by the group. To determine whether individual variability is spatially heterogeneous across the cortex, individual variability values were plotted on the cortical surface and statistically thresholded with a jack-knife procedure (*p <* 0.05), separately for each participant (see Methods). This procedure retained only voxels with extreme individual variability values. Finally, to identify regions with low or high individual variability at the group level, thresholded participant-specific maps were projected to the template surface *fsaverage* (Fischl et al., 1999) and averaged across all participants.

Figure 3 shows the group-averaged individual variability map, along with participant-specific maps for two representative individuals (Supplementary Figure 4 shows all participants). These maps reveal that the largest individual differences in lexical-semantic representations are found in the temporo-parietal junction (TPJ), angular gyrus, Precuneus, ventral portions of the superior temporal sulcus (STS), and patches in the frontal cortex. Previous research suggests that these areas function as convergence zones, integrating modal and multimodal information to support conceptual knowledge (Binder & Desai, 2011; Fairhall & Caramazza, 2013; Pulvermüller, 2013). The smallest individual differences are found in the dorsal bank of the STS, in the ventral temporal lobe, and in portions of ventral Precuneus and prefrontal cortex. These regions with low individual variability are close to sensory areas such as primary auditory cortex, or they are predominantly tuned to concrete information such as visual-related concepts (e.g., voxels colored in green in Figure 1). Taken together, these results demonstrate that individual differences in lexical-semantic representations are spatially heterogeneous across the cortex. The most variable representations are found in higher-order associative areas within the distributed conceptual network. The least variable representations are found in lexical-semantic areas located near primary perceptual areas.

**Figure 3.**
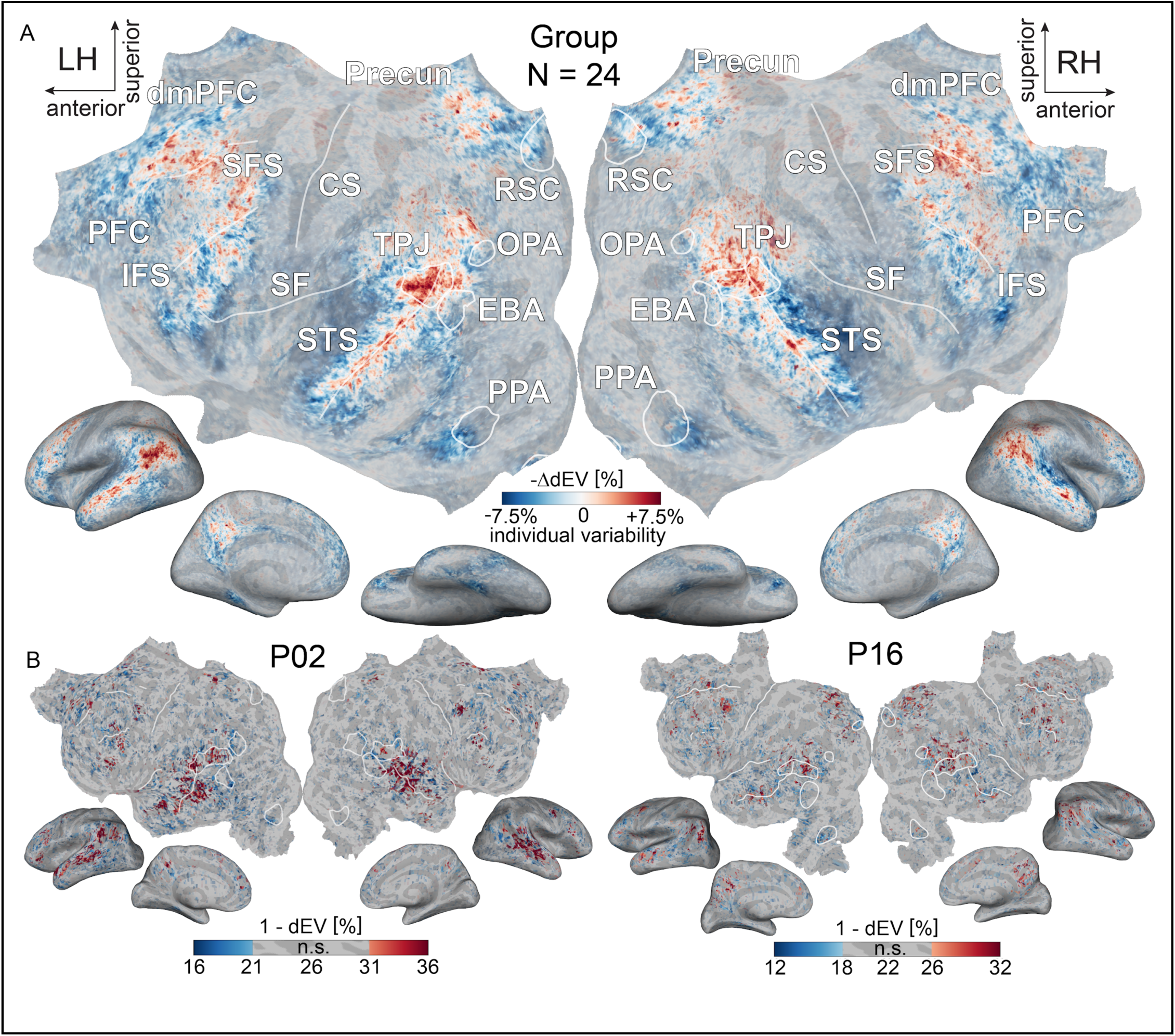
Spatial heterogeneity of individual differences in lexical-semantic representations. Prior work suggests that individual differences in brain anatomy and function are spatially heterogeneous. However, whether this heterogeneity extends to complex lexical-semantic representations remains unknown. To address this question, we modified the dEV summary statistic to measure individual variability in each voxel. Because the dEV quantifies the variance explained by the group, individual variability is expressed as 1 - dEV. Individual variability maps were created separately for each participant and statistically thresholded at p < 0.05 using a jack-knife procedure (see Methods). [A] Group-averaged individual variability map (N=24) plotted on the template surface fsaverage. Blue indicates low individual variability; red indicates high individual variability. Values are expressed as relative change from the average. [B] Participant-specific individual variability maps for two representative participants (P02 and P16; other participants are shown in Supplementary Figure 4). These maps show that largest individual differences are found in the TPJ, Precuneus, STS, and parts of the dorsomedial prefrontal cortex. These regions with high individual variability are thought to function as convergence zones, integrating modal and multimodal information to support conceptual knowledge. The smallest individual differences are found in the dorsal bank of the STS, in the ventral temporal lobe, and in portions of ventral Precuneus and prefrontal cortex. These regions with low individual variability are close to sensory areas such as primary auditory cortex, or they are predominantly tuned to concrete information such as visual-related concepts. These results demonstrate that associative and high-order areas exhibit the largest individual differences in lexical-semantic representations.

### Representations of social-related concepts show the largest individual differences

The previous analysis demonstrated that associative and high-order cognitive areas show the largest individual differences in lexical-semantic representations. The lexical-semantic maps in Figure 1 suggest that these regions predominantly represent social concepts. This pattern led us to hypothesize that representations of social-related information may show higher individual variability than representations of other types of information.

Testing this hypothesis requires comparing individual variability values between brain regions that represent social concepts versus other concepts. However, the narrative stories evoke hundreds of different lexical-semantic concepts, making it impractical to compare individual variability for each concept separately. To address this challenge, a data-driven procedure called model connectivity was used to cluster encoding model weights into functional networks with distinct selectivity (Meschke et al., 2023; see Methods). In the story-listening experiment used here, model connectivity recovers 12 networks that are broadly distributed across the cortex and that generalize across participants (Meschke et al., 2023). Each network represents a distinct category of lexical-semantic concepts. Six networks represent social- and discourse-related concepts (plotted in shades of red and pink in Figure 4). The remaining six networks represent physical- and place-related concepts (plotted in shades of green in Figure 4). Together, these 12 networks span the entire lexical-semantic space (Figure 4). If our hypothesis is correct, individual variability should concentrate in networks representing social concepts rather than physical concepts. To test this prediction, each participant’s total individual variability was partitioned across the 12 lexical-semantic categories defined by the networks. Each voxel was assigned to the category for which that voxel showed the strongest tuning. Individual variability for each category was computed by averaging across voxels assigned to that category. These values were then expressed as fractions of each participant’s total individual variability to account for different numbers of voxels across participants. Finally, these fractions were averaged across all participants. Statistical significance was assessed by comparing the observed fractions against chance level, determined by random permutations across voxels (FDR-corrected non-parametric permutation test, see Methods).

**Figure 4.**
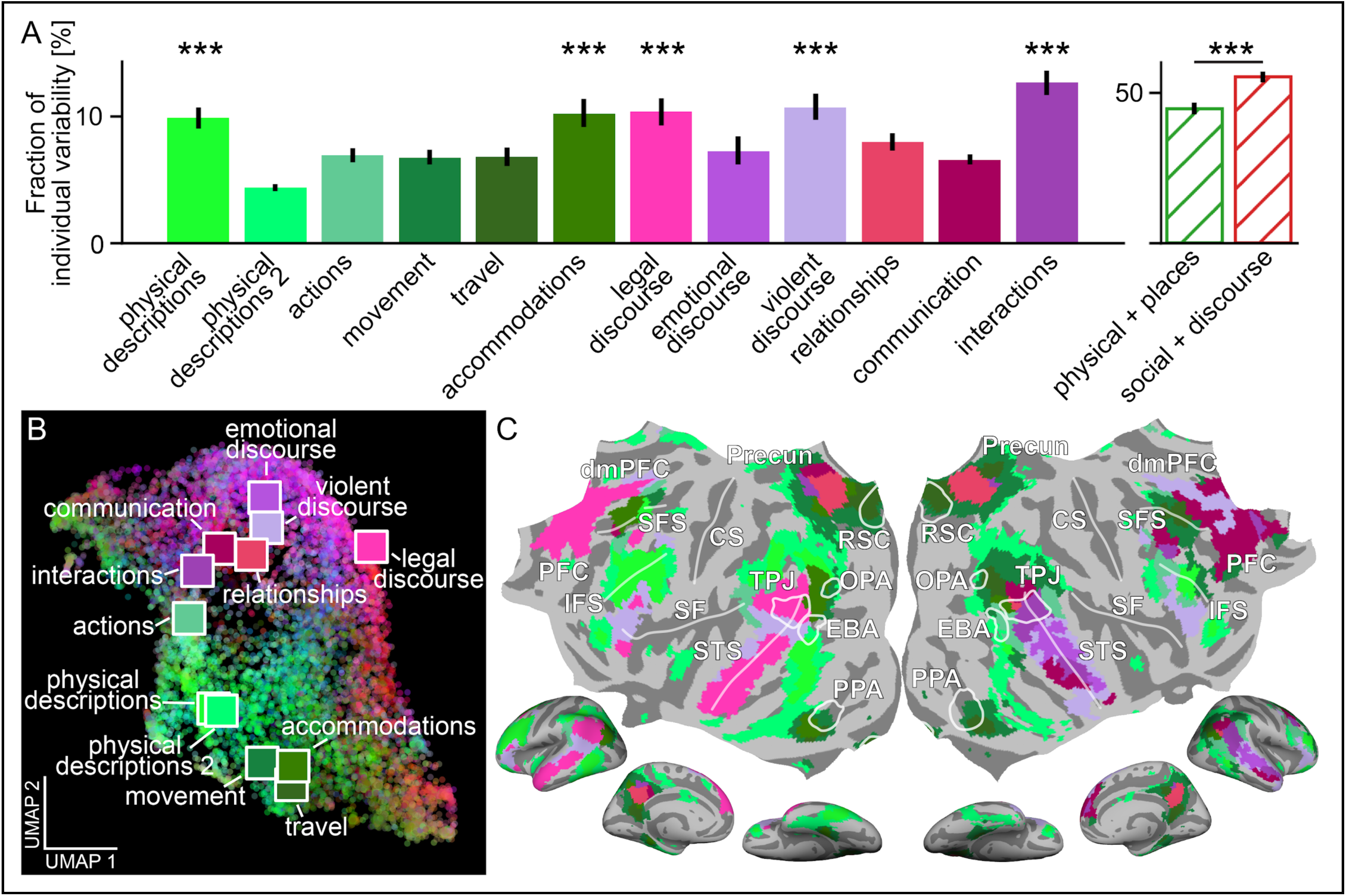
Representations of social concepts show the largest individual differences. Individual variability in lexical-semantic representations concentrates in associative and high-order cognitive areas that predominantly represent social concepts (Figures 1 and 3). This pattern led us to hypothesize that representations of social-related concepts may show higher individual variability than representations of other types of concepts. To test this hypothesis, a data-driven clustering approach called model connectivity was used to cluster lexical-semantic model weights into 12 networks with distinct functional selectivity. Each network represents a distinct category of lexical-semantic concepts. Six networks represent social- and discourse-related concepts (colored in shades of red/pink), while six networks represent physical- and place-related concepts (colored in shades of green). [A] Individual variability was partitioned across the 12 categories and expressed as fractions of total individual variability for each participant. The barplot shows the group-averaged fractions across participants (error bars indicate 95% bootstrapped confidence intervals). Five categories show significantly greater individual variability than chance (*p* < 0.001, FDR-corrected non-parametric permutation test): three social categories (*interactions*, *violent discourse*, *legal discourse*) and two physical categories (*physical descriptions*, *accommodations*). The right panel compares broad categories by grouping social concepts (red) versus physical concepts (green). Social concepts account for 55.31% of individual variability compared to 44.69% for physical concepts, with a significant difference of 10.61% (*p <* 0.001, non-parametric permutation test). [B] The 985-dimensional lexical-semantic space is visualized by projecting the entire feature space onto a reduced 2D space, obtained with the Uniform Manifold Approximation and Projection (UMAP) dimensionality reduction algorithm. Each dot indicates a word from the *english1000* vocabulary, colored by the RGB scheme from Figure 1. Colored squares mark category locations. This visualization shows that the 12 categories defined by the model connectivity networks span the major dimensions of the lexical-semantic space. [C] Cortical distribution of the 12 lexical-semantic networks identified by model connectivity. Networks selective to social-related concepts overlap with areas showing the largest individual variability. These results demonstrate that representations of social concepts exhibit significantly greater individual variability than representations of physical concepts.

The results indicate that five of the 12 major lexical-semantic categories show significantly greater individual variability than chance (Figure 4). Three categories with significant individual variability are related to social- and discourse-related concepts: social interactions (associated with words such as *friend*, *calls*; fraction of individual variability = 12.61%, 95% CI: 11.67% to 13.59%, *p* < 0.001, FDR-corrected non-parametric permutation test), violent discourse (*killing, stabbed*; 10.67%, 95% CI: 9.67% to 11.77%, *p* < 0.001), and legal discourse (*judge, innocent*; 10.34%, 95% CI: 9.25% to 11.40%, *p* < 0.001). The remaining two categories with significant individual variability are related to physical- and place-related concepts: physical descriptions (*colored*, *inches*; 9.84%, 95% CI: 9.00% to 10.68%, *p* < 0.001) and accommodations (*bedroom*, *apartment*; 10.16%, 95% CI: 9.09% to 11.31%, *p* < 0.001). To determine whether social-related representations exhibit the largest individual differences, in a post-hoc analysis the 12 lexical-semantic categories were grouped into two broad categories of social and physical concepts (Figure 4). Across all 24 participants the social category accounts for 55.31% of individual variability (95% CI: 53.42% to 57.19%), while the physical category accounts for 44.69% (95% CI: 42.81% to 46.58%), with a significant difference of 10.61% (95% CI: 6.87% to 14.41%, *p* < 0.001). Thus, the social category accounts for significantly more individual variability compared to the physical category. These results confirm that representations of social concepts exhibit greater individual variability than representations of physical concepts.

### Individual differences in lexical-semantic representations reflect person-specific conceptual biases

The results presented thus far demonstrate clear individual differences in conceptual representations. Here we present a method for interpreting these individual differences. Recall that in the framework used in this study, individual differences are quantified by the difference between each participant’s lexical-semantic model weights and the group (Figure 2). These weight differences reveal systematic biases towards specific lexical-semantic concepts. For example, if one person’s brain responds to social concepts more strongly compared to the group, their weight differences will point toward social representations in the lexical-semantic space. Similarly, if another person’s brain responds more strongly to spatial and temporal concepts, their weight differences will point toward spatial and temporal representations. These systematic biases reflect how each person’s conceptual space differs from the conceptual space of others.

To identify the major dimensions underlying the conceptual biases of each participant, principal component analysis (PCA) was applied to the weight differences separately for each participant. This approach reduces the high-dimensional weight differences into a smaller set of interpretable dimensions that capture the major axes of individual variation. Each resulting principal component (PC) reflects a specific dimension in lexical-semantic space along which one participant differs systematically from the group. To ensure that the PCs reflected true individual differences rather than noise, a bootstrapping procedure was used to quantify the reliability of each PC, and to retain only the most reliable components for each participant (see Methods). Across the 24 participants, the median number of PCs required to reliably capture individual differences is 18.5 (min: 9, max: 26). These PCs explain on average 75.63% of the individual variability in each participant (95% CI: 73.77% to 77.40%). Thus, compared to the size of the lexical-semantic space (985 dimensions), a relatively small number of lexical-semantic dimensions captures most individual differences in conceptual representations.

To demonstrate how these participant-specific dimensions can be interpreted, we focus on the first two PCs of each participant. These two dimensions capture the largest sources of individual variability for each participant. However, the same interpretation procedure can be easily applied to all reliable PCs to obtain fine-grained descriptions of individual differences. Figure 5 shows the first two PCs for three representative participants (P02, P06, and P22), and Supplementary Figures 5-8 show all participants. The first two dimensions of each participant explain on average 37.19% of total individual variability within the same participant (95% CI: 35.72% to 38.73%), but explain only 1.44% of individual variability in other participants (95% CI: 1.34% to 1.55%). This specificity demonstrates that the dimensions underlying each participant’s conceptual biases are unique to that person.

**Figure 5.**
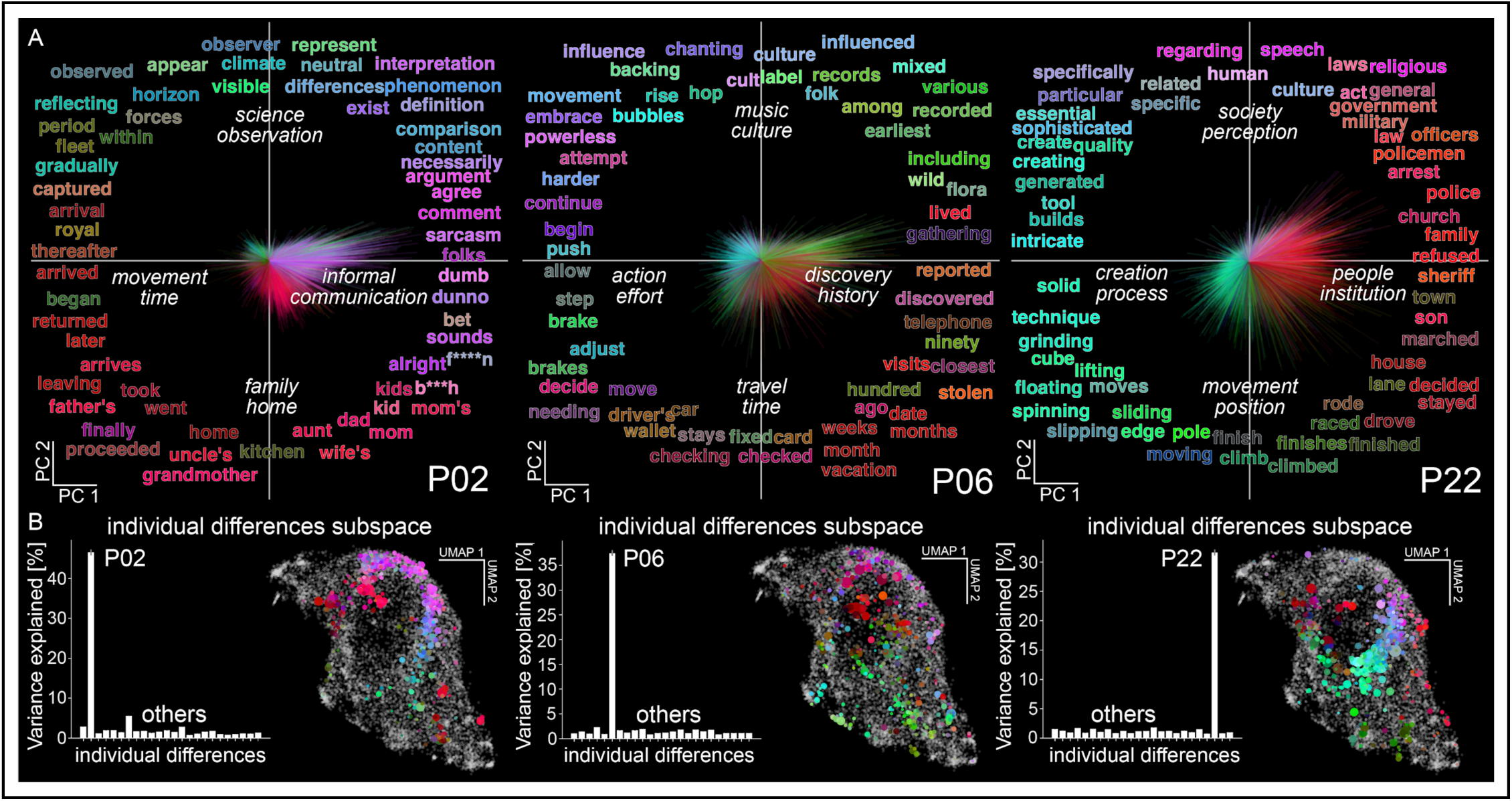
Individual differences in lexical-semantic representations reflect person-specific conceptual biases. In the framework used here, individual differences are quantified using optimal transport to measure how each participant’s lexical-semantic model weights differ from the group distribution of model weights. These differences reveal systematic biases towards specific lexical-semantic concepts, reflecting how each person’s conceptual space differs from others. To identify the major dimensions underlying these person-specific conceptual biases, principal component analysis was applied separately to the weight differences of each participant. [A] The first two principal components (PCs) for three representative participants (P02, P06, P22) are shown as orthogonal axes. (Supplementary Figures 5-8 shows all other participants.) Words from the english1000 vocabulary that project strongly onto each dimension are displayed and colored according to the RGB color scheme from Figure 1. Lines correspond to individual voxel biases towards specific lexical-semantic concepts. The line length is proportional to bias magnitude, and the color matches the closest word. White labels in italics are generated by an automated algorithm based on large language models, and describe the concepts associated with each PC direction (see Methods). This analysis reveals systematic biases for P02 towards concepts related to informal communication, for P06 towards discovery and history, and for P22 towards people and institutions. [B] The bar plots show the fraction of individual variability explained by the first two PCs. The first two dimensions of each participant explain more than 30% of total individual variability within the same participant, but less than 2% of individual variability in other participants. This specificity demonstrates that the dimensions underlying each participant’s conceptual biases are unique to that person. The UMAP visualization highlights the words in the english1000 vocabulary (colored dots) with the largest projection onto the subspace spanned by the first two PCs. This visualization demonstrates that the subspaces of different participants span distinct portions of the lexical-semantic space, confirming that these individuals differ along fundamentally different conceptual axes. Taken together, these results reveal that conceptual representations differ between individuals along lexical-semantic dimensions that are uniquely associated with each person. These person-specific dimensions reflect systematic biases in how different individuals represent conceptual information.

To interpret the concepts reflected by these dimensions, we used a data-driven interpretation algorithm (Antonello et al., 2024; Singh et al., 2023). This automated algorithm uses a large language model (LLM) to generate multiple concept descriptions from words in the *english1000* vocabulary that project strongly onto each dimension, then selects the best description based on a quantitative score (see Methods). Figure 5A displays the resulting descriptions for the three representative participants (Supplementary Tables 1, 2 list interpretations for all participants). These descriptions reveal distinct concepts associated with each participant. In participant P02 the first dimension separates concepts related to *movement* and *time* from those related to *informal communication*. The second dimension separates concepts related to *family* and *home* from those related to *science* and *observation*. In participant P06, the first dimension separates concepts related to *action* and *effort* from those related to *discovery* and *history*. The second dimension separates concepts related to *travel* and *time* from those related to *music* and *culture*. In participant P22, the first dimension separates concepts related to *creation* and *process* from those related to *people* and *institutions*. The second dimension separates concepts related to *movement* and *position* from those related to *society* and *perception*. For each participant, these two dimensions define a two-dimensional subspace within the full lexical-semantic space. Figure 5B shows a UMAP visualization of these participant-specific subspaces (Supplementary Figures 5-8 shows all participants). This plot reveals that the subspaces of different participants span distinct portions of the lexical-semantic space, confirming that these individuals differ along fundamentally different conceptual axes. Taken together, the results described here reveal that conceptual representations differ between individuals along lexical-semantic dimensions that are uniquely associated with each person. These person-specific dimensions reflect systematic biases in how different individuals represent conceptual information.

## Discussion

The study of individual differences in brain function is critical for distinguishing natural variations from the impact of cognitive syndromes and disorders. However, current neuroscience and clinical neurology studies predominantly focus on group-level effects, discarding the critical information required to understand individual differences. To overcome the limitations of group-level studies, here we developed a new statistical framework for measuring and interpreting individual differences in complex functional brain representations. We applied this framework to examine conceptual representations evoked by narrative stories. Our results reveal clear individual differences in how people represent conceptual information during story-listening. These individual differences manifest as systematic biases towards specific lexical-semantic concepts. These biases do not reflect canonical semantic dimensions such as abstract/concrete (Binder et al., 2009; Hoffman & Bair, 2024; Montefinese, 2019; Tang et al., 2021; Wang et al., 2010), animate/inanimate (Caramazza & Shelton, 1998; Connolly et al., 2012; Huth et al., 2012; Sha et al., 2014), or social/nonsocial (Diveica et al., 2024; Han et al., 2024; Haxby et al., 2020; Rice et al., 2018) (see Supplementary Results and Supplementary Figure 9). Rather, these biases are organized along lexical-semantic dimensions that are unique to each participant. These idiosyncratic lexical-semantic dimensions provide an interpretable fingerprint of how each individual represents conceptual information evoked by the narrative stories.

We suspect that the conceptual biases that we report here reflect individual variations in cognitive traits and personal life experiences. This hypothesis is supported by the spatial distribution of individual variability (Figures 2 and 3). Our analysis revealed that the TPJ, Precuneus, STS, and patches in the frontal cortex exhibit the largest individual differences. Previous work shows that these areas function as multimodal convergence zones (Binder & Desai, 2011; Fairhall & Caramazza, 2013; Pulvermüller, 2013), combining information from different modalities to support conceptual knowledge and its integration with long-term memory (Gallant & Popham, 2020). Recent theories posit that these areas also integrate ongoing sensory information with a person’s prior beliefs, memories, and experiences (Fernandino & Binder, 2024; Yeshurun et al., 2021). Therefore, lexical-semantic representations in these areas may depend on personal experience and cognitive traits. This interpretation is consistent with our finding that representations in these areas are the most variable across individuals.

While an explanation of individual variability in terms of personal experience and cognitive traits is compelling, alternative explanations are possible. Because our framework relies on encoding models, the observed individual differences could reflect trivial between-participant differences in model performance, or in demographics such as age or gender. However, none of these factors significantly predict the observed individual differences (see Supplementary Results). Another concern is that the measured individual differences reflect variations in cognitive states, arousal, or attention during story listening, rather than stable cognitive traits. However, this explanation is highly unlikely for several reasons. Encoding models were estimated on two hours of data, collected over multiple scanning runs and sessions for each participant. This extensive sampling of brain activity effectively averages out fluctuations in cognitive states, arousal, or attention. For the encoding models to reflect such fluctuations, these effects would need to persist systematically across different days. By definition, such sustained effects would reflect cognitive traits rather than states. Therefore, the findings reported here likely reflect individual differences in conceptual representation that stem from personal experience and cognitive traits.

The individual differences framework developed here overcomes the limitations of prior research, and paves the way for new research directions. Previous approaches have focused either on functional networks with limited interpretability (Cui et al., 2020; Finn & Bandettini, 2021; Gordon et al., 2015; Gratton et al., 2018; Mueller et al., 2013; Vanderwal et al., 2017), or on low-dimensional representations that fail to capture the complexity of brain function (Botch & Finn, 2024; Chadwick et al., 2016; Charest et al., 2014; Grabner et al., 2007; Rypma & D’Esposito, 1999). Our framework offers a general, quantitative approach to systematically measure and characterize individual differences in functional representations. Because this framework is based on voxelwise encoding models, functional representations can be recovered from a wide range of experimental paradigms, including complex naturalistic tasks. By combining precision high-dimensional functional brain mapping with extensive behavioral assays, this approach reveals how brain function mediates complex human behavior in individual people.

Our framework also provides new opportunities for developing fMRI applications to study, diagnose, and monitor cognitive and developmental disorders. Current diagnostic approaches for conditions such as Alzheimer’s disease, frontotemporal dementia, schizophrenia, or autism spectrum disorder rely predominantly on behavioral, anatomical, or genetic markers. However, these markers have significant limitations. Behavioral deficits often overlap across disorders. Anatomical markers may be undetectable in vivo. Clear genetic markers remain unavailable for many disorders. Our individual differences framework can address these limitations by measuring and interpreting differences in functional representations caused by these conditions. By comparing functional representations in individual patients against normative control groups, this framework can reveal the dimensions that are most diagnostic for each disorder. These dimensions could produce precise, patient-specific functional brain markers that complement existing diagnostics.

## Methods

### Participants

Twenty-four participants (11 females and 13 males) between the ages of 21 and 35 took part in the experiment. Nineteen participants were scanned at the UC Berkeley Brain Imaging Center, and five participants were scanned at the UT Austin Biomedical Imaging Center. The data from these latter five participants are a subset of the public dataset by LeBel et al. (2023). The public dataset includes two additional participants (UTS06 and UTS08) who were excluded from the current analyses due to excessive motion (more than 10% of volumes with a framewise displacement > 0.5 mm). The public dataset also includes another participant (UTS04) who was not included in the current analyses because of missing data (fMRI responses to the *life* story are not available for this participant). All participants were healthy with normal hearing. All the experimental procedures were approved by the UC Berkeley Committee for Protection of Human Subjects or by the Institutional Review Board at the University of Texas at Austin. All participants provided written informed consent to participate in the experiments.

### Experimental procedure

Participants listened to eleven 10- to 15-min stories taken from *The Moth Radio Hour* over multiple scanning sessions (see Huth et al., 2016 and LeBel et al., 2023 for more information about this paradigm). Participants were instructed to close their eyes and to listen to the stories. Auditory stimulation was delivered over Sensimetrics S14 in-ear piezoelectric headphones (Sensimetrics Corporation, Malden, MA). For the participants at UC Berkeley, a Behringer Ultra-Curve Pro hardware parametric equalizer was used to flatten the frequency response of the headphones based on calibration data provided by Sensimetrics. For the participants at UT Austin, the audio files were filtered via software based on calibration data provided by Sensimetrics (see LeBel et al., 2023 for more information). Sound volume was adjusted for each participant at the beginning of every scanning session.

### MRI data collection

For the 19 participants at UC Berkeley, MRI data were collected on a 3T Siemens TIM Trio scanner with a 32-channel Siemens volume coil at the UC Berkeley Brain Imaging Center. For 18 of the 19 participants, gradient-echo EPI images were collected with a sequence that had the following parameters: repetition time (TR) = 2.0045 s, echo time (TE) = 35 ms, flip angle = 74°, voxel size = 2.24 × 2.24 × 4.13 mm (slice thickness = 3.5 mm with 18% slice gap), matrix size = 100 × 100, field of view = 224 × 224 mm, anterior-posterior phase encoding direction. A custom-modified bipolar water-excitation radiofrequency pulse was used to avoid signal from fat. For the remaining participant (P05), gradient-echo EPI images were collected with a multi-band sequence that had the following parameters: repetition time (TR) = 1.16 s, echo time (TE) = 34 ms, flip angle = 62°, multi-band factor (simultaneous multi-slice) = 3, voxel size = 2.5 mm x 2.5 mm x 2.5 mm (slice thickness = 2.5 mm), matrix size = 84 x 84, field of view = 210 x 210 mm, anterior-posterior phase encoding direction. The field of view covered the entire cortex for all participants. For 16 of 19 participants, a gradient-echo fieldmap scan was acquired at the beginning of each functional run for EPI distortion correction. For two of the remaining three participants (P05, P15), two spin-echo volumes with opposite phase-encoding direction were acquired at the beginning of each functional run for EPI distortion correction. For the remaining participant (P20), fieldmap scans were not collected. Up to four anatomical T1-weighted multi-echo MP-RAGE sequences were collected for each participant. These sequences used the parameters recommended by FreeSurfer for optimizing surface segmentation.

For the five participants at UT Austin (P07, P08, P10, P11, P12), MRI data were collected on a 3T Siemens Skyra scanner with a 64-channel Siemens volume coil at the UT Austin Biomedical Imaging Center. Gradient-echo EPI images were collected with a multi-band sequence that had the following parameters: repetition time (TR) = 2.00 s, echo time (TE) = 30.8 ms, flip angle = 71°, multi-band factor (simultaneous multi-slice) = 2, voxel size = 2.6 mm x 2.6 mm x 2.6 mm (slice thickness = 2.6 mm), matrix size = 84 x 84, field of view = 220 x 220 mm, anterior-posterior phase encoding direction. The field of view covered the entire cortex for all participants. No fieldmap scans were collected for these participants. An anatomical T1-weighted multi-echo MP-RAGE sequence was collected for each participant. This sequence used the parameters recommended by FreeSurfer for optimizing surface segmentation. For more information about the data from these participants, we refer the reader to the original paper describing the dataset (LeBel et al., 2023).

Despite different collection sites and fMRI sequences in the two datasets, no apparent group differences emerged in the experimental results. Model prediction accuracy maps, lexical-semantic maps, and individual variability maps showed no systematic differences between the two participant groups (see Supplementary Results and Figures). Therefore, all analyses presented in the main text combine data from both datasets.

### Data preprocessing

FreeSurfer (Fischl, 2012) was used to generate high-resolution surface reconstructions from the T1-weighted scan. For each participant, surface reconstructions were manually inspected and adjusted. Relaxation cuts were performed manually to obtain a flatmap version of the surface.

Each functional run was motion-corrected, distortion-corrected (if fieldmap scans were available), and aligned to the participant’s anatomical volume. Motion correction transformations were computed with the FMRIB Linear Image Registration Tool (FLIRT) from FSL (Jenkinson et al., 2002; Jenkinson & Smith, 2001). For each run, a reference volume was computed by temporally averaging the motion-corrected volumes. This reference volume was used as the moving volume for each run to compute additional transformations. If gradient-echo fieldmap scans were available, distortion correction transformations were computed with FUGUE from FSL (Smith et al., 2004). If spin-echo volumes with opposite phase-encoding direction were available, fieldmap estimates were computed with TOPUP from FSL (Andersson et al., 2003; Smith et al., 2004) and distortion correction transformations were computed with FUGUE from FSL. If fieldmap scans were not available, the preprocessing pipeline did not include distortion correction. Anatomical alignment transformations were computed with *bbregister* from FreeSurfer (Greve & Fischl, 2009). Finally, ANTs with Lanczos interpolation was used to combine and apply all transformations in a single interpolation step (Avants et al., 2009). For all participants, the aligned and temporally averaged volumes of each run were inspected visually to check the alignment to the participant’s anatomical volume. The resulting preprocessed data were then temporally detrended and denoised by including six noise components estimated with CompCor (Behzadi et al., 2007). For the participant (P05) with fMRI data sampled at a TR of 1.16 s, after detrending and denoising *scipy.signal.interpolate* was used to temporally resample their data to a 2s TR. Prior to model estimation the preprocessed data were temporally z-scored separately within each run. To ensure that only gray matter voxels were included in the analyses, after preprocessing the functional data were masked with Pycortex’s *line_nearest* mask (Gao et al., 2015). This mask selects only voxels that overlap with the cortical ribbon, which is defined by the pial and white matter surfaces generated by FreeSurfer.

The preprocessing pipeline was implemented in Python, and it included custom Python code and functions from nipype (Esteban et al., 2020; Gorgolewski et al., 2011). Note that this pipeline differs from the pipeline used in many of the previous publications from our laboratory. This new pipeline adopts several improvements first developed and validated by fMRIPrep (Esteban et al., 2019), such as using *bbregister* and ANTs to align functional images to each participant’s anatomical volume.

### Voxelwise encoding models

Voxelwise encoding models use features extracted from the experimental stimulus to predict brain responses in each participant. Here, lexical-semantic features and low-level speech features were extracted from the narrative stories, as described below. Regularized linear regression was used to predict brain responses simultaneously from the lexical-semantic and low-level features. Separate training and test sets were used to estimate and evaluate voxelwise encoding models. For each participant, the training set consisted of 10 stories (3737 samples, ∼124 minutes of data) and the test set consisted of one story repeated between two and four times (291 samples for each stimulus repetition, ∼10 minutes of data). All voxelwise encoding models were estimated and evaluated in each participant separately.

### Feature spaces

The lexical-semantic content of the narrative stories was quantified by extracting lexical-semantic features from the stimuli. The lexical-semantic features were based on a 985-dimensional word-embedding space, the *english1000* feature space (see Huth et al., 2016 for more details on this lexical-semantic feature space). Each word from the transcript of the stories was projected onto the *english1000* feature space. The lexical-semantic vectors were then resampled at times corresponding to the fMRI acquisitions using a three-lobe Lanczos filter with the cutoff frequency set to the Nyquist frequency of the fMRI acquisition (0.25 Hz for a 2s TR).

To take into account low-level speech information that may correlate with lexical-semantic information, a low-level speech feature space was created by extracting multiple nuisance features. The low-level speech feature space had 41 total features and included 39 phonemic features that capture how often each of the phonemes in English was spoken over time; a single feature that counts the number of phonemes spoken over time (phoneme-rate); and a single feature that counts the number of words spoken over time (word-rate) (Deniz et al., 2019).

### Feature preprocessing

Before model estimation, each stimulus feature was z-scored over time within each run. To account for the hemodynamic delay, voxelwise encoding models were estimated with a Finite Impulse Response (FIR) model that included five temporal delays (1, 2, 3, 4, and 5 TRs). This was accomplished by concatenating feature vectors that had been delayed by 1, 2, 3, 4, and 5 TRs (in this dataset corresponding to time delays of 2 s, 4 s, 6 s, 8 s, and 10 s). Taking the dot product of this concatenated feature space with a set of linear weights is equivalent to a convolution with linear temporal kernels that have non-zero entries for 1, 2, 3, 4, and 5 TRs.

### Model estimation

Models were estimated independently for each participant. A modified version of ridge regression called banded ridge regression was used to model the functional data (Dupré la Tour et al., 2022; Nunez-Elizalde et al., 2019). Banded ridge regression implements a separate regularization hyperparameter for each feature space. Two separate regularization hyperparameters were optimized: one for the lexical-semantic feature space and one for the low-level speech feature space. The regularization hyperparameters were optimized independently for each voxel.

The optimal regularization hyperparameters were determined via cross-validation. The entire training set of *n* functional runs was split into a smaller training set and a validation set. The smaller training set consisted of *n-1* runs, and the validation set consisted of one run (leave-one-run-out cross-validation scheme). Then, multiple banded ridge regression models were fit to the smaller training set with 20 logarithmically spaced regularization hyperparameters ranging from 10^1^ to 10^20^. The prediction accuracy of each of these models was estimated on the validation set. This process was repeated by leaving out each of the *n* training runs. The resulting prediction accuracies were then averaged across left-out validation runs, and the hyperparameters with the highest prediction accuracy were selected separately for each voxel.

All model fitting was performed using *himalaya*, a Python package developed by our lab to efficiently fit ridge and banded ridge regression models on CPU and GPU (Dupré la Tour et al., 2022).

### Model evaluation

The prediction accuracy of the final voxelwise encoding models was evaluated on a held-out test set. For each voxel, the coefficient of determination *R*^2^ was used to quantify the prediction accuracy of the model that included all feature spaces. The prediction accuracy of each separate feature space was quantified with the split coefficient of determination 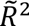. The split coefficient of determination decomposes the overall model prediction accuracy to reveal the individual contribution of each feature space (Dupré la Tour et al., 2022). In this study the model prediction accuracy is decomposed into the contributions of the lexical-semantic feature space and the low-level speech feature space, that is 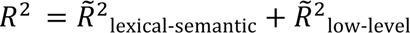.

### Model weight adjustments

Before interpreting the voxelwise encoding model weights, the weights need to be adjusted to account for potential differences across voxels. These differences are caused by differences in hemodynamic responses, noise levels, and regularization hyperparameters.

#### Averaging model weights across delays

Because each voxelwise encoding model includes a finite impulse response (FIR) model, each stimulus feature is distributed across five temporal delays (as described above in *Feature preprocessing*). Therefore, after model estimation each voxel is associated with five model weights for each feature. These five model weights correspond to the five temporal delays of the FIR model. To collapse across the five temporal delays, the five model weights for each feature were averaged across delays. As a result of this procedure, each voxel is associated with a 985-dimensional vector of lexical-semantic model weights.

#### Re-scaling model weights

To optimize model prediction accuracy within each voxel, the regularization hyperparameter is determined separately for each voxel. Differences in signal-to-noise ratio (SNR) between voxels can cause them to have different regularization hyperparameters. Because the regularization hyperparameters directly affect the scale of the estimated model weights, differences in scale across voxels may reflect differences in SNR rather than differences in functional representations. As a consequence, any difference in scale across voxels caused by the regularization hyperparameters can bias the interpretation of the model weights.

One solution to address this bias is to use a global regularization hyperparameter that is optimized across all voxels (Huth et al., 2016). Using a global regularization hyperparameter ensures that all weights have a similar scale, but because the hyperparameter is not optimized separately for each voxel, this approach leads to worse model prediction accuracy. Therefore, to maximize model prediction accuracy we optimized the regularization hyperparameters separately for each voxel (see *Model estimation* above). To prevent biases caused by differences in scale of the model weights, we implemented an additional rescaling step. The lexical-semantic model weights in each voxel were first scaled to the unit norm, and then they were re-scaled by the cross-validated split prediction accuracy in the training set (negative prediction accuracies were set to zero).

Mathematically, let 𝑤 ∈ ℝ^985^ be the lexical-semantic model weights in a voxel and 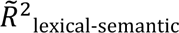 the split prediction accuracy of the lexical-semantic feature space in that voxel. The re-scaled weight is

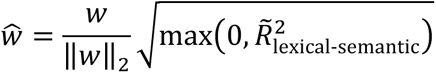

We found empirically that this re-scaling step corrects for differences in the scale of the model weights while maximizing model prediction accuracy within each voxel (Dupré la Tour et al., 2025; Meschke et al., 2023).

#### Participant-specific cortical maps of lexical-semantic tuning

The lexical-semantic model weights of a voxel describe the tuning of that voxel for multiple lexical-semantic features. However, because these model weights are high-dimensional (985 lexical-semantic features), it is impractical to create cortical maps that accurately reflect the full dimensionality of the fit voxelwise models. Therefore, to create cortical maps that describe the lexical-semantic tuning it is first necessary to perform some form of dimensionality reduction. Here we projected the high-dimensional lexical-semantic model weights onto a reduced 3-dimensional space that was obtained from a previous study (Huth et al., 2016). That earlier study produced this reduced lexical-semantic space by performing principal component analysis (PCA) on the lexical-semantic model weights of a group of six participants, and then retaining the first three principal components (PCs) for visualization.

Here, for each individual participant cortical maps of lexical-semantic tuning were created by coloring each voxel according to the projection of that voxel’s model weights onto the reduced 3D lexical-semantic space. That is, the 3D projection vector is considered as a vector in the RGB color space. Each voxel is thus colored according to the resulting RGB value. With this visualization, different colors correspond to representations of different lexical-semantic concepts. To highlight only well predicted areas, the opacity of each voxel was set to be proportional to the model prediction accuracy in the training set. To do this, the cross-validated prediction accuracy in the training set was clipped between 0 and a robust maximum (corresponding to the 95th percentile across all voxels with positive prediction accuracy), and then it was rescaled between 0 and 1. The resulting value was used as the opacity value.

We chose this visualization approach over statistical thresholding because the cross-validated prediction accuracy in the training set provides a continuous measure of effect size computed on independent validation sets. It is easy to interpret and does not require an arbitrary threshold such as a p-value, which could hide voxels with low but reliable prediction accuracy.

### Group-level cortical map of lexical-semantic tuning

To create a group-level cortical map of lexical-semantic tuning, the participant-specific voxelwise lexical-semantic weights were first projected to the template surface *fsaverage* (Fischl et al., 1999). This projection was computed by combining two separate transformations: a volume-to-surface transformation and a surface-to-template transformation. The first transformation projects each participant’s volumetric data to their own cortical surface. Pycortex’s *line_nearest* mapper was used to resample data from voxels to vertices (Gao et al., 2015). For each vertex, this mapper samples the functional data at 64 evenly spaced intervals between the inner (white matter) and outer (pial) surfaces of the cortex. Samples are taken according to nearest-neighbor interpolation, and each sample is given the value of its enclosing voxel. The resulting samples are then averaged and assigned to the vertex. The second transformation projects each participant’s surface data to the *fsaverage* template surface. Pycortex’s *get_mri_surf2surf_matrix* was used to compute the surface-to-template projection matrix from FreeSurfer’s *mri_surf2surf*. To reduce the propagation of numerical errors, the volume-to-surface and surface-to-template projection matrices were combined into a single volume-to-template projection matrix. For each participant, this surface-to-template projection matrix was applied to the voxelwise lexical-semantic model weights to obtain vertexwise model weights in *fsaverage* space. The resulting model weights in *fsaverage* space were averaged across participants and projected onto the reduced 3D lexical-semantic space. Finally, the same visualization procedure described above for participant-specific cortical maps was used to produce the group-level cortical map of lexical-semantic tuning.

#### Quantifying functional individual differences as differences in distributions of encoding model weights

The lexical-semantic model weights of a voxel describe the tuning of that voxel for multiple lexical-semantic features. The model weights are estimated with banded ridge regression, which imposes an L2 regularization on the model weights. L2 regularization is equivalent to imposing a Gaussian prior on the model weights. Therefore, the set of all lexical-semantic model weights can be modeled as a multivariate Gaussian distribution in the *english1000* lexical-semantic feature space. For each participant, the distribution of lexical-semantic model weights determines which lexical-semantic features are represented in that participant’s brain activity during the experiment. Differences in distributions of lexical-semantic model weights between individuals correspond to differences in lexical-semantic representation. Under this formulation, the problem of quantifying functional individual differences is equivalent to quantifying differences in high-dimensional distributions of encoding model weights.

To describe possible approaches for comparing distributions of voxelwise encoding model weights, we consider the more general problem of comparing two distributions of samples in a high-dimensional space. We refer to these distributions as the *source* distribution and the *target* distribution. (For example, a source distribution could be the distribution of one participant’s model weights and the target distribution could be the distribution of the model weights from an independent group of participants.) Mathematically, we denote 𝑊_𝑠_ ∈ ℝ^𝑝×𝑢^ the *u* source samples and 𝑊_𝑡_ ∈ ℝ^𝑝×𝑣^ the *v* target samples, where *p* is the number of dimensions and *p < u*, *p < v*. In the context of voxelwise encoding models, *p* is the dimensionality of the feature space, while *u* and *v* are the number of voxels.

Because these distributions are Gaussian, they can be summarized by their mean and covariance. For simplicity, we assume that both 𝑊_𝑠_, 𝑊_𝑡_ have a zero mean, that is they are centered along the second dimension. The sample covariance of the source distribution is given by 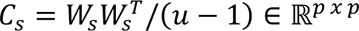, and the sample covariance of the target distribution is given by 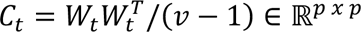. To simplify the notation, in the following paragraphs we will omit the normalization by (𝑢 − 1) or (𝑣 − 1) when describing the sample covariance. Note that this corresponds to a simple change of variable 𝑊′_𝑠_ = 𝑊_𝑠_/√𝑢 − 1. Additionally, we will assume that both 𝐶_𝑠_, 𝐶_𝑡_ are full-rank, or equivalently that all their eigenvalues are non-zero.

Principal Component Analysis (PCA) is a common approach for comparing high-dimensional distributions of samples. PCA identifies the principal components (PCs) of a distribution, ordered by decreasing explained variance. To compare two distributions, one can quantify the fraction of variance in the source distribution that is explained when the distribution is projected onto a subset of the target PCs. If this value is sufficiently large, then it could be concluded that the two distributions are similar. However, this approach has two key limitations. First, the projection of the source samples onto the target PCs ignores the eigenvalues of each PC (i.e., the amount of variance the PCs explain in the target distribution). Second, if the source distribution is projected onto all target PCs, then the entire source variance is explained. Therefore, one would mistakenly conclude that the two distributions are identical. The Supplementary Methods provide detailed mathematical derivations of these limitations. Because of these limitations, a different statistical approach is needed to appropriately compare two distributions.

### The distributional Explained Variance (dEV)

To accurately compare two distributions of samples, it is necessary to preserve all the information contained in both distributions. Because a zero-centered Gaussian distribution can be entirely summarized by its covariance, a measure that compares covariances would take into account the entire information contained in both distributions.

Several measures for comparing covariances already exist (Fletcher & Joshi, 2007; Peyré & Cuturi, 2019). However, these measures have an important drawback for fMRI applications. Because they operate only on covariances, they discard any information associated with individual samples. In the context of voxelwise encoding models, each sample is a model weight from a specific voxel. Thus, these measures would discard the information provided by the anatomical location of the voxel. This information is crucial for interpreting neuroscience results based on fMRI data. To solve this problem, here we developed a novel measure that can be used to compare distributions of samples while preserving information associated with individual samples. This novel measure is based on the mathematical theory of the optimal transport (Peyré & Cuturi, 2019; Villani, 2008), and we call it the distributional Explained Variance (dEV).

To derive the dEV, we first observe that the projection of 𝑊_𝑠_ on its first *k* PCs can be also defined as the transform 𝑇(𝑊_𝑠_) that minimizes the low-rank reconstruction error of 𝑊_𝑠_(Hastie et al., 2009). This transform is defined as

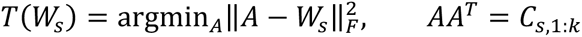

where ‖⋅‖_𝐹_ is the Frobenius norm, and 𝐶_𝑠,1:𝑘_ is the covariance matrix 𝐶_𝑠_ truncated after the *k* largest eigenvalues. That is, 𝐶_𝑠,1:𝑘_ = 𝑈𝛬^2^_1:𝑘_𝑈^𝑇^, where 𝑈 ∈ ℝ^p×p^ is an orthonormal matrix, and 𝚲_1:k_ ∈ ℝ^p×p^ is a diagonal matrix filled with singular values 𝜆_𝑖_ such that 𝜆_1_ ≥ 𝜆_2_ ≥ ⋯ ≥ 𝜆_𝑘_ > 0, and 𝜆_𝑖_ = 0, 𝑖 = 𝑘 + 1, …, 𝑝.

This optimization problem can be modified to compare two arbitrary distributions of samples. We seek a transformation *T* of the source samples such that the transformed source samples are minimally modified, and their covariance is equal to the covariance of the target samples. The transformation corresponds to the solution of the optimization problem

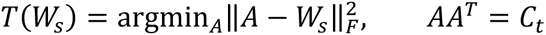

where now 𝐶_𝑡_ is the covariance of the target samples. We refer to 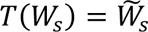 as the *transformed* source samples. When this transformation is applied to voxelwise lexical-semantic model weights, the individual weights are transformed in the high-dimensional lexical-semantic feature space, but their anatomical location is preserved.

A solution to this optimization problem is the optimal transport transform. For Gaussian distributions, the optimal transport transform can be derived analytically in closed form (Peyré & Cuturi, 2019). In this case, the solution corresponds to 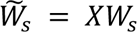, where

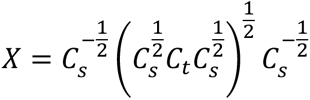

is the optimal transport transform for Gaussian distributions.

Following the logic of PCA to compare the source and target distributions, the total variance of 𝑊_𝑠_ can be compared to the variance explained by 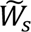. Because 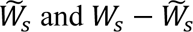 are not always orthogonal, it is not possible to use a strict definition of explained variance as in the case of PCA (see Supplementary Methods). Therefore, we need to relax the notion of explained variance in the same way that it is relaxed to compute the coefficient of determination *R^2^* in regression problems. In regression, *R^2^* is defined as 1 − 𝑆𝑆_res_/𝑆𝑆_tot_, where 𝑆𝑆_res_ is the residual sum of squares and 𝑆𝑆_tot_ is the total sum of squares. Similarly, the dEV can be expressed as

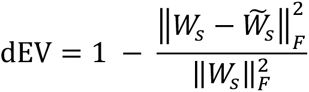

The dEV is interpreted analogously to the coefficient of determination *R^2^*. dEV values range from] − ∞, 1], with larger values corresponding to a greater similarity between the distribution of source samples and the distribution of target samples.

The dEV generalizes the concept of explained variance of a subspace spanned by the first *k* PCs. Indeed, if the target covariance equals the truncated covariance 𝐶_𝑠,1:𝑘_, then the dEV equals the fraction of variance explained by the first *k* PCs. To see why, consider that when 𝐶_𝑡_ = 𝐶_𝑠,1:𝑘_, the optimal transport transform is given by

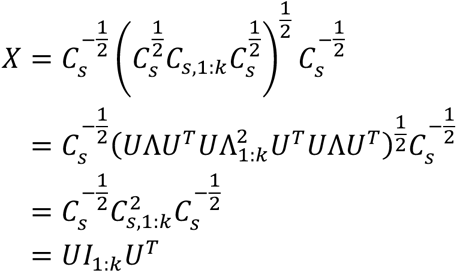

where 𝐼_1:𝑘_ ∈ ℝ^p×p^ is diagonal matrix such that 𝐼_(𝑖𝑖)_ = 1, 𝑖 = 1, …, 𝑘 and 0 otherwise. With this transform, the transformed source samples are given by

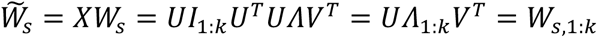

where 𝑊_𝑠,1:𝑘_ is the projection of 𝑊_𝑠_ on the subspace spanned by the first *k* PCs. In this case, the dEV equals to

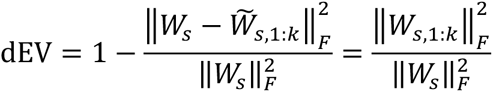

which equals to the fraction of variance explained by the first *k* PCs (see Supplementary Methods).

#### Using the dEV to quantify functional individual differences in lexical-semantic representations

The dEV can be used to examine individual differences in high-dimensional distributions of lexical-semantic model weights. For a group of *N* participants, we use a leave-one-subject-out procedure to compare each participant’s distribution of model weights to the group distribution consisting of the weights from the other *N-1* participants. For one of these iterations, the source samples are the model weights from one participant and the target samples are the model weights from all other participants. To group weights across participants, the weights are concatenated along the voxel dimension. The dEV is then used to quantify the similarity of that participant to the independent group of participants.

### Jack-knife confidence intervals for the dEV values

A block jack-knife procedure was used to estimate the variability of the dEV values for each participant. In this procedure, new banded ridge regression models were fit on all possible sets of *n-1* training runs for each participant. The model weights were estimated by fixing the optimal hyperparameters found by cross-validation in the training set. (This is equivalent to storing the model weights associated with the optimal hyperparameters in each cross-validation training fold during the hyperparameter search.) This procedure generated a set of *n-1* lexical-semantic model weights for each voxel and each participant. These lexical-semantic model weights were adjusted as described above (*Model weights adjustments*) and were used to compute a set of *n-1* jack-knifed dEV values for each participant. Finally, these jack-knifed dEV values were used to estimate the jack-knife variance 𝜎_𝑑𝐸𝑉_ and a 95% confidence interval according to dEV ± 𝑡_0.25,𝑛–1_ ⋅ 𝜎_dEV_, where 𝑡_0.25,𝑛–1_ is the 0.25-level critical value of a Student’s *t* distribution with *n-1* degrees of freedom (Abdi & Williams, 2010).

### Estimates of individual variability in each voxel

To estimate the variability of functional representations in each voxel, a dEV value was first derived for each voxel. This was accomplished by changing the Frobenius norm to the L2 norm over the first dimension of 𝑊_𝑠_. Therefore, given a set of model weights 𝑤_𝑖_ ∈ ℝ^985^ in a voxel *i*, its dEV value is defined as

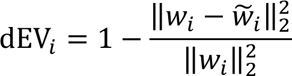

where ‖⋅‖_2_ is the L2 norm. For each participant, these voxelwise dEV values were statistically thresholded at *p* < 0.05, using a global test based on the jack-knife confidence intervals. That is, the dEV value of a voxel was declared significantly different from the global dEV value if dEV_𝑖_ < dEV − 𝑡_0.25,𝑛–1_ ⋅ σ_dEV_ or dEV_𝑖_ > dEV + 𝑡_0.25,𝑛–1_ ⋅ σ_dEV_. The individual variability in each statistically significant voxel was then defined as the amount of variance unexplained by the group, that is 1 − dEV_𝑖_.

A group-averaged map was created by first expressing the individual variability in each voxel as the relative change from the global dEV value for each participant. That is, each statistically significant voxel was assigned the value (1 − dEV_𝑖_) − (1 − dEV) = −(dEV_𝑖_ − dEV) = −ΔdEV. Then, the resulting statistically thresholded maps were projected to the *fsaverage* template surface (Fischl et al., 1999). The projection followed the same process described in the previous section *Group-level cortical map of lexical-semantic tuning*. After projection of the individual participants’ maps, a weighted average was used to compute a single group-level map across participants. For each participant, the value in a vertex was weighted by the cross-validated training prediction accuracy. This was done to ensure that the group map would not be biased by participants whose models had overall low prediction accuracy.

In the surface visualization, the opacity of each vertex was set to be proportional to the model prediction accuracy in the training set. First, the model prediction accuracy was averaged across participants (arithmetic average). Then, the resulting values were clipped between 0 and a robust maximum (corresponding to the 95th percentile across all voxels with positive prediction accuracy). Finally, these values were rescaled between 0 and 1 and used as the opacity value for each vertex. We used this visualization to highlight the areas that contributed the most to the dEV value. Prior to computing the dEV value in a voxel, the model weights in that voxel are rescaled by the model prediction accuracy (see the section *Model weight adjustments*). Therefore, each voxel contributes to the dEV measure proportionally to its model prediction accuracy. In the group map, this contribution is reflected by the opacity of the vertex.

#### Estimates of individual variability across different lexical-semantic categories

To quantify individual variability across lexical-semantic categories, we first assigned each voxel to one of 12 major lexical-semantic categories based on its lexical-semantic model weights. These categories were derived from a previous study that introduced model connectivity—a data-driven approach that recovers functional networks with consistent tuning from voxelwise encoding model weights (Meschke et al., 2023). In that study, model connectivity was applied to fMRI data from nine participants listening to the same narrative stories used here. (Eight of those participants are included in the current study.)

Model connectivity identified 12 lexical-semantic networks with consistent tuning, and one noise network comprising brain areas not activated by the story-listening task. Each lexical-semantic network is associated with a 985-dimensional network-average weight vector in the *english1000* feature space. To determine the lexical-semantic category represented in each voxel, these network-average weight vectors were used as cluster centroids. For each voxel, we computed the cosine similarity between the lexical-semantic model weights in that voxel and the network-averaged model weights, and assigned that voxel to the lexical-semantic category with the highest similarity. This process partitioned each participant’s voxels into 12 lexical-semantic clusters. To account for varying cluster sizes, the individual variability was averaged across all voxels within each cluster. Finally, these averages were divided by the sum over all 12 clusters to express them as fractions of total individual variability.

To assess statistical significance, non-parametric permutation testing was used to generate a null distribution and compute empirical p-values. Null distributions for each participant were created by randomly permuting individual variability estimates across voxels while maintaining the voxel assignments to the 12 lexical-semantic categories (9999 permutations). A group-level null distribution was created by averaging the participants’ null distributions. The resulting p-values were corrected for multiple comparisons using the Benjamini-Hochberg false discovery rate (FDR) procedure (Benjamini & Hochberg, 1995).

### UMAP visualization of the word-embedding lexical-semantic space

To visualize the entire word-embedding space and the relationship between words, the Uniform Manifold Approximation and Projection (UMAP) dimensionality reduction algorithm was used to project the 985-dimensional word vectors onto a two-dimensional space. The UMAP algorithm was implemented in the Python package *umap* (McInnes et al., 2018). To improve the resulting visualization, the parameters of the algorithm were manually changed to use 100 neighbors (n_neighbors=100) and a cosine distance metric with a minimum distance of 0 (metric=“cosine”, min_dist=0). Because the UMAP algorithm is stochastic, a fixed random seed was selected to ensure replicability of the visualization. Note that different UMAP parameters will produce different two-dimensional embeddings. However, this is irrelevant because we used UMAP for visualization purposes only, and we did not interpret the precise distances between points in the two-dimensional space.

### Recovering idiosyncratic lexical-semantic subspaces for each participant

Although PCA is not ideal for quantifying the similarity of distributions, it is an excellent approach for recovering the major dimensions and subspaces underlying a single distribution. Thus, to recover the unique lexical-semantic subspaces associated with each participant, PCA was first performed separately for each participant on the distribution of weight differences 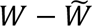. (Recall that 𝑊 is the participant’s lexical-semantic model weights and 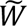is the transformed model weights.) PCA produces a set of orthogonal dimensions, ordered according to the amount of variance explained by each dimension. Some of these dimensions reliably capture individual differences, while other dimensions capture noise. To focus on signal dimensions, a bootstrapping procedure was used to determine the reliable PCs. For each bootstrap iteration, voxels were sampled with replacement, and PCA was performed again on the bootstrapped sample of weight differences. Then, for each of the original PCs the maximum absolute correlation was computed between that PC and all bootstrapped PCs. This procedure was repeated for 1,000 iterations to generate a distribution of correlation values for each original PCs. Finally, the reliability of each PC was quantified as the median of the bootstrap distribution (Lebart et al., 1995). The first consecutive PCs with a reliability of at least *r* = 0.9 were retained for further analysis.

#### Summarizing idiosyncratic lexical-semantic dimensions for each participant

To summarize the concepts associated with each participant’s idiosyncratic lexical-semantic dimension, we first projected all the 10,470 words in the vocabulary of the *english1000* word-embedding space onto each idiosyncratic dimension. To increase interpretability, we removed 419 words that did not provide lexical meaning but that were nonetheless present in the vocabulary (e.g., articles and determiners, pronouns, conjunctions, prepositions, auxiliary verbs, etc.). The remaining 10,051 words were then sorted according to their projection on each lexical-semantic dimension. The 50 words with the smallest projection and the 50 words with the largest projection were selected to identify the lexical-semantic concepts associated with the extremes of each dimension.

An iterative procedure called Summarize And Score (SASC; Antonello et al., 2024; Singh et al., 2023) was used to summarize each set of words into a few labels. As the name suggests, this procedure consists of two steps: summarization and scoring. In the summarization step, a random subset of 40 words was selected and GPT-4o (OpenAI) was prompted to provide a summary that best explained the set of 40 words. In the scoring step, GPT-4o was asked to generate a list of 20 words associated with the summary (positive examples) and 20 words not associated with the summary (negative examples). For any words generated by GPT-4o that were absent from the *english1000* vocabulary, the *word2vec* embedding space (Mikolov et al., 2013) was used to identify their closest equivalents within the *english1000* vocabulary. The positive and negative example words were then projected on the lexical-semantic dimension being summarized, and a score was computed by taking the difference between the average projection of the positive example words and the average projection of the negative example words. This procedure was repeated five times, and the summary with the highest score was finally selected. The output of GPT-4o was minimally modified to parse the response.

The GPT-4o prompt used for the summarization step was:

*Here is a list of words. Focus on the meaning of the words. Summarize their common concept into a short list of words, using the lowest number of words sufficient to summarize the list*.

*Use one word if one word is sufficient, and use two words if two words are sufficient, and so on. You can use up to four words*.

*Do not use words that describe only grammar, syntax, or part of speech, such as “adverbs”, “adjectives”, “nouns”, etc*.

*Explain your reasoning and provide some example words in the list when giving your explanation*.

*Always answer in the following format: “The common concept among these words is: [word_1, word_2, …]*.

*The concept is [word_1] because [your explanation]*.

*The concept is [word_2] because [your explanation]*.

...

*Here is a list of words:*

*{words}*

*The common concept among these words is:*

The GTP-4o prompt used for the scoring step was:

*Generate 20 words that are {blank_or_not}related to the concept of {concept}. 1*.

### Data analysis implementation

All the analyses reported here were implemented in Python with the following packages: numpy (van der Walt et al., 2011), scipy (Virtanen et al., 2020), scikit-learn (Abraham et al., 2014), nipype (Esteban et al., 2020; Gorgolewski et al., 2011), nibabel (Brett et al., 2024), pyRiemann (Barachant et al., 2025), pycortex (Gao et al., 2015), cottoncandy (Nunez-Elizalde et al., 2018), himalaya (Dupré la Tour et al., 2022), pytorch (Paszke et al., 2019), cupy (Okuta et al., 2017), imodelsX (https://github.com/csinva/imodelsX), and matplotlib (Hunter, 2007).

### Data and materials availability

A repository with the data and code used in this study are being prepared for public release. It will be made available upon publication.

