## Supplementary Materials for "Individual differences shape conceptual representation in the brain"

### Supplementary Results

#### Individual differences are not explained by differences in prediction accuracy or demographics

Figure 2D in the Main Text reveals that dEV values vary across individuals, ranging from a minimum of 73.87% to a maximum of 80.74%. This variability indicates that some participants have lexical-semantic representations that are less similar to the group (low dEV), while others have representations that are more similar to the group (high dEV). However, these variations could also stem from trivial variability across participants, such as differences in model prediction accuracy or demographics. To rule out this possibility, we examined the relationship between dEV values and model prediction accuracy, participant age, and participant sex.

Prediction accuracy does not explain individual differences. To determine whether differences in prediction accuracy could explain the observed variability in dEV, we correlated median training and test prediction accuracy with dEV values across all participants. Because the test dataset is created by averaging across repetitions of the same stimulus (see Methods) and participants had different numbers of repetitions (min 2, max 4), for this analysis the test model prediction accuracy was computed on the average of only two stimulus repetitions for all participants. Training prediction accuracy showed no significant correlation with dEV values: $r = 0.06$, 95% CI: -0.36 to 0.51 (bootstrap CI), $p = 0.778$ (non-parametric correlation permutation test). Test prediction accuracy also showed no significant correlation: $r = 0.06$, 95% CI: -0.26 to 0.44, $p = 0.774$. These results indicate that differences in model prediction accuracy do not explain the observed variability in dEV values.

Demographics do not explain individual differences. To determine whether differences in participant age could explain the observed variability in dEV, participant age was correlated with dEV values. This correlation was not statistically significant: $r = 0.19$, 95% CI: -0.21 to 0.54, $p = 0.365$. To determine whether participant sex could explain the observed variability in dEV, dEV values were compared between male (N=13) and female participants (N=11). The average dEV value for male participants was 77.34% (95% CI: 76.24% to 78.40%), while the average dEV value for female participants was 78.08% (95% CI: 77.29% to 78.91%). The difference was not statistically significant: -0.74%, 95% CI: -2.11% to 0.57%, $p = 0.337$ (two-independent-samples non-parametric permutation testing). These results indicate that differences in demographics do not explain the observed variability in dEV values.

The results from these control analyses indicate that variability in dEV values reflect genuine individual differences in how participants represent conceptual information, rather than methodological confounds or demographic factors.

#### Effects of model performance on the spatial heterogeneity of individual differences

The analyses in Figure 3 in the Main Text reveal that individual differences in lexical-semantic representations are spatially heterogeneous across the cortex. Because model prediction accuracy is also spatially heterogeneous (see Supplementary Figure 1), spatial differences in individual variability could be explained by spatial differences in model prediction accuracy. To address this concern, we correlated individual variability values (1 - dEV) with model prediction accuracy across all voxels for each participant.

Across the 24 participants, individual variability showed a very weak but statistically significant positive correlation with training prediction accuracy: $r = 0.079$, 95% CI: 0.045 to 0.109, $p < 0.001$ (one-sample non-parametric permutation testing). Individual variability also correlated very weakly with test prediction accuracy: $r = 0.076$, 95% CI: 0.041 to 0.108, $p < 0.001$. These positive correlations indicate that larger individual differences tend to occur in regions with higher model prediction accuracy. However, the magnitude of the correlations indicates that this effect is quite small; model prediction accuracy explains only $r^2 = \sim 0.6\%$ of the spatial distribution of individual variability. Thus, even though the correlation is statistically significant, model prediction accuracy does not meaningfully account for the observed spatial heterogeneity of individual differences.

**Canonical semantic axes do not explain participant-specific conceptual biases**

Previous research suggests that conceptual knowledge is organized across several core semantic axes, including social/nonsocial (Diveica et al., 2024; Han et al., 2024; Haxby et al., 2020; Rice et al., 2018), animate/inanimate (Caramazza & Shelton, 1998; Connolly et al., 2012; Huth et al., 2012; Sha et al., 2014), abstract/concrete (Binder et al., 2009; Hoffman & Bair, 2024; Montefinese, 2019; Tang et al., 2021; Wang et al., 2010), and positive/negative valence (Lettieri et al., 2019; Lindquist et al., 2016). Because of their importance in the literature, we asked whether these four canonical semantic axes explain the participant-specific conceptual biases revealed by this study (Figure 5 in the Main Text).

The four canonical semantic axes were constructed by selecting words from the *english1000* vocabulary that reflect the extremes of each semantic dimension. To identify these words, established normed lexicons provided ratings for socialness (Diveica et al., 2023), animacy (VanArsdall & Blunt, 2022), concreteness (Brysbaert et al., 2014), and valence (Mohammad & Turney, 2013). For each dimension, the 40 highest-rated and 40 lowest-rated words were selected to reflect the two extremes (complete word sets are provided in Supplementary Table 3). For each extreme, a vector was generated by averaging the *english1000* word vectors corresponding to that extreme and normalizing to unit length. The final semantic axis was created by computing the normalized difference between the two extreme vectors. For example, the social/nonsocial axis was constructed by selecting the word vectors associated with 40 highest-rated and the 40 lowest-rated words on socialness. These word vectors were averaged separately for each extreme to create a "social" vector and a "nonsocial" vector. To create the social/nonsocial dimension, the "nonsocial" vector was subtracted from the "social" vector.

To assess whether these canonical semantic axes could explain the participant-specific conceptual biases, we calculated the fraction of variance explained by each axis. Supplementary Figure 9 shows that across all 24 participants each canonical semantic axis explains less than 2% of the variance in the conceptual biases. By contrast, the first two participant-specific lexical-semantic dimensions explain on average more than 35% of the variance in the participant-specific conceptual biases (Supplementary Figure 9). These results indicate that canonical semantic axes do not explain the participant-specific conceptual biases observed during narrative story listening. Instead, these biases reflect unique, participant-specific combinations of lexical-semantic concepts.

### Supplementary Methods

#### Noise ceiling correction

The estimates of model prediction accuracy are affected by the level of noise in each voxel. Because the noise level varies across brain areas and participants, raw prediction accuracy values cannot be compared fairly across brain areas or participants. To account for differences in noise level, the raw prediction accuracy values on the held-out test dataset are typically normalized by the noise ceiling of each voxel (Hsu et al., 2004; Sahani & Linden, 2002; Schoppe et al., 2016). The noise ceiling corresponds to the maximum prediction accuracy that can be obtained in a voxel, given the measured test dataset. To compute the noise ceiling, it is useful to first compute a related quantity called the explainable variance (EV) of the measured signal. Let $Y_k$ be the signal measured in a voxel over the *k*-th repetition of the test stimulus (k = 1, …, N) and let $\tilde{Y} = \frac{1}{N}\sum_{i=1}^{N} Y_k$ be the average over the repetitions of the test stimulus. An unbiased estimator of the explainable variance is given by

$$\mathrm{EV} = \frac{1}{N-1}\left[N\,\mathrm{Var}(\tilde{Y}) - \frac{1}{N}\sum_{i=1}^{N}\mathrm{Var}(Y_i)\right]$$

where Var indicates the variance computed over time samples. As the name suggests, EV reflects the maximum amount of variance in the measured signal that can be explained by a model. Finally, the noise ceiling $R^2_{\mathrm{max}}$ is given by

$$R^2_{\mathrm{max}} = \frac{\mathrm{EV}}{\mathrm{Var}(\tilde{\mathrm{Y}})}$$

To account for differences in noise levels, the noise ceiling is used to normalize the raw prediction accuracy. That is, for each voxel the normalized prediction accuracy is equal to

$$R^2_{\mathrm{norm}} = \frac{R^2}{R^2_{\mathrm{max}}}$$

However, for very noisy voxels the estimated noise ceiling $R^2_{\mathrm{max}}$ can be lower than the measured prediction accuracy $R^2$. In these voxels, the normalized prediction accuracy diverges to values greater than 1. To correct for this divergence, we first identified the set of voxels for which the noise ceiling is lower or equal than the model prediction accuracy, that is all voxels for which $R^2_{\mathrm{max}} \leq R^2$. We then selected the maximum $R^2_{\mathrm{max}}$ over this set of voxels. Finally, we set the $R^2_{\mathrm{max}}$ of all voxels to be above this maximum value. This conservative procedure ensures that the noise ceiling is greater than the measured prediction accuracy in all voxels.

#### Statistical thresholding of prediction accuracy maps

Non-parametric permutation testing was used to statistically threshold the participant-specific normalized test prediction accuracy maps, displayed in Supplementary Figures 1 and 2. To generate a null distribution of prediction accuracy values on the test set, the joint predictions of the banded ridge regression model were randomly permuted over time in each voxel. The joint prediction corresponds to the prediction of the lexical-semantic feature space plus the prediction of the low-level speech feature space (see Methods in Main Text). As in previous works (Chen et al., 2024; Deniz et al., 2019), blocks of 10 temporally contiguous samples (corresponding to 20 s of data) were randomly permuted to account for the existing temporal autocorrelation in the data. Then, the permuted joint prediction accuracy was calculated. This process was repeated 9,999 times to generate a null distribution. An empirical p-value was calculated for each voxel by counting the number of times the permuted prediction accuracy was greater than the non-permuted prediction accuracy. This number was divided by the total number of permutations to obtain an empirical p-value, adding 1 to both the numerator and denominator to avoid p-values equal to 0 (Phipson & Smyth, 2010). To account for multiple tests, the empirical p-values were corrected with the Benjamini-Hochberg false-discovery rate procedure (Benjamini & Hochberg, 1995). Finally, a mask of statistically significant voxels was generated by thresholding the FDR-corrected p-value map at $p < 0.05$. The same mask was applied to the maps for the lexical-semantic and low-level speech feature spaces. To threshold the group-averaged prediction accuracy map, a group-level union mask was created by combining each participant-specific significance mask projected to the *fsaverage* template surface. This union mask was additionally thresholded to highlight only vertices that were significant for at least 25% of participants (N=6).

**Limitations of Principal Component Analysis (PCA) to compare distributions of samples**

As we describe in the Methods section of the Main Text, we seek an approach to compare a source distribution of model weights $W_s \in \mathbb{R}^{p\times u}$ to a target distribution of model weights $W_t \in \mathbb{R}^{p\times v}$. The most common approach for comparing high-dimensional distributions of samples is Principal Component Analysis (PCA). PCA recovers the directions of largest variance in a distribution of samples. These directions are referred to as the *principal components* (PCs). To compare two distributions of samples, a straightforward approach is to compare the variance explained by their respective PCs. If the PCs of the target distribution explain sufficiently high variance in the source distribution, it could be concluded that the two distributions are similar. However, this intuitive approach has several drawbacks. To see why, it is useful to first describe mathematically how PCA is computed, and how it is used.

An efficient way to compute PCA on a set of samples is through singular value decomposition (SVD). With SVD, $W_s = U\Lambda V^T$, where $U \in \mathbb{R}^{p\times p}$ is an orthonormal matrix, $\Lambda \in \mathbb{R}^{p\times p}$ is a diagonal matrix filled with the singular values $\lambda_k$ such that $\lambda_1 \geq \lambda_2 \geq \ldots \geq \lambda_p > 0$, and $V \in \mathbb{R}^{p\times p}$ is an orthonormal matrix. Given the SVD of $W_s$, its covariance is equal to $C_s = W_s {W_s}^T = U\Lambda V^T V\Lambda U^T = U\Lambda^2 U^T$. Therefore, the singular values $\lambda_k$are equal to the square root of the eigenvalues of the covariance matrix. This equality shows that PCA can be computed both by SVD of the centered samples or by the eigen-decomposition of the covariance matrix of the samples. By rearranging the SVD of $W_s$, we obtain $U^T W_s = \Lambda V^T$. The column vectors of $\Lambda V^T$ are the *principal components (PCs)* of $W_s$, and the orthonormal matrix $U$ projects the samples $W_s$ onto these PCs.

<u>Explained variance of principal components.</u> The principal components of a distribution of samples can be examined in terms of the amount of variance that they explain. To determine the amount of variance explained by one PC, let $U_k \in \mathbb{R}^{p\times 1}$ be the *k*-th column of $U$. The projection of $W_s$ on the *k*-th PC is

$$U^T{}_k W_s = U^T{}_k U\Lambda V^T = \lambda_k V^T$$

and its variance equals

$$\|\lambda_k V^T\|_F^2 = \lambda_k^2$$

Where $\|\cdot\|_F$ indicates the Frobenius norm. The variance associated with the *k*-th PC is referred to as the *explained variance* of the *k*-th PC.

When multiple PCs are considered together, they span a subspace of the original sample space. To determine the amount of variance explained by such a subspace, let use denote $W_{s,1:k} = U_{1:k}\Lambda V^T$ the projection of $W_s$ onto the subspace spanned by the first *k* PCs, where $U_{1:k} \in \mathbb{R}^{p\times p}$ is a matrix containing the first *k* columns of $U$ and 0 otherwise. The total explained variance of the first *k* PCs is

$$\left\|W_{s,1:k}\right\|_F^2 = \|U_{1:k}\Lambda V^T\|_F^2 = \mathrm{tr}\left(U_{1:k}\Lambda \mathrm{V}^\mathrm{T}\mathrm{V}\Lambda \mathrm{U}_{1:\mathrm{k}}^T\right) = \sum_{i=1}^{k} \lambda_i^2$$

where the second equality follows from the definition of the Frobenius norm, and $\mathrm{tr}(\cdot)$ denotes the trace of a square matrix. Therefore, the explained variance of the subspace spanned by the first *k* PCs is equal to the sum of the explained variance of each of the first *k* PCs. Note that because $U, V$ are orthonormal, the projection of the samples onto the first *k* PCs $W_{s,1:k}$ and the residuals $W_s - W_{s,1:k}$ are orthogonal. Therefore, the total variance of $W_s$ can be decomposed as

$$\|W_s\|_F^2 = \left\|W_{s,1:k}\right\|_F^2 + \left\|W_s - W_{s,1:k}\right\|_F^2$$

This equality shows that the decomposition of the total variance into the variance explained by single or multiple PCs is a valid variance decomposition.

<u>Explained variance of a subspace.</u> The explained variance of a subspace is an intuitive concept that is not limited to the PCs of the source samples $W_s$. For example, the source samples $W_s$ can be projected onto the subspace spanned by the first *k* PCs of the target samples $W_t$. Then, one can compare the variance of the source samples explained by the first *k* target PCs to the variance explained by the first *k* source PCs. If these two quantities are sufficiently similar, it can be concluded that the first *k* PCs of both the source samples $W_s$ and the target samples $W_t$ span a similar subspace. This would be evidence that the source and target distribution are similar, at least within a lower-dimensional subspace.

To see how this approach works, let $W_t = U'\Lambda'V'^T$ be the SVD of the target samples $W_t$. The projection of $W_s$ onto the first *k* PCs of $W_t$ is given by $U'^T_{1:k}W_s$, and the variance explained by these first *k* target PCs is

$$\left\|U'^T_{1:k}W_s\right\|_F^2 = \mathrm{tr}(U'^T_{1:k}W_sW_s^TU'_{1:k}) = \mathrm{tr}(U'^T_{1:k}U\Lambda^2U^TU'^T_{1:k}\ )$$

Given a fixed number of PCs $k < p$, this projection can be used to compare the explained variance of the first *k* source PCs of $W_s$ to the explained variance of the first *k* target PCs of $W_t$. That is, $\left\|W_{s,1:k}\right\|_F^2 = \|U^T_{1:k}W_s\|_F^2$ is compared to $\|U'^T_{1:k}W_s\|_F^2$.

This approach is simple and intuitive, but it has two serious limitations. First, the projection of the source samples onto the target PCs ignores the singular values $\Lambda'$ of the target samples $W_t$. (Recall that the singular values $\Lambda'$ are the square root of the eigenvalues of $C_t = W_tW_t^{\ T}$.) Therefore, this projection discards the important information provided by the covariance of the target samples $W_t$, and it only compares the PCs of $W_s$ to the principal *directions* of $W_t$. Second, if all the PCs of the target samples $W_t$ are retained, the projection will explain the entire variance in the source samples. In this case,

$$\|U'^TW_s\|_F^2 = \mathrm{tr}(U'^TW_sW_s^TU') = \mathrm{tr}(W_sW_s^T) = \|W_s\|_F^2$$

where the second equality follows from the cyclic properties of the trace. Therefore, when all target PCs are retained, the projection is independent of the source and target PCs, and the source and target distributions cannot be compared appropriately.

### Supplementary Figures

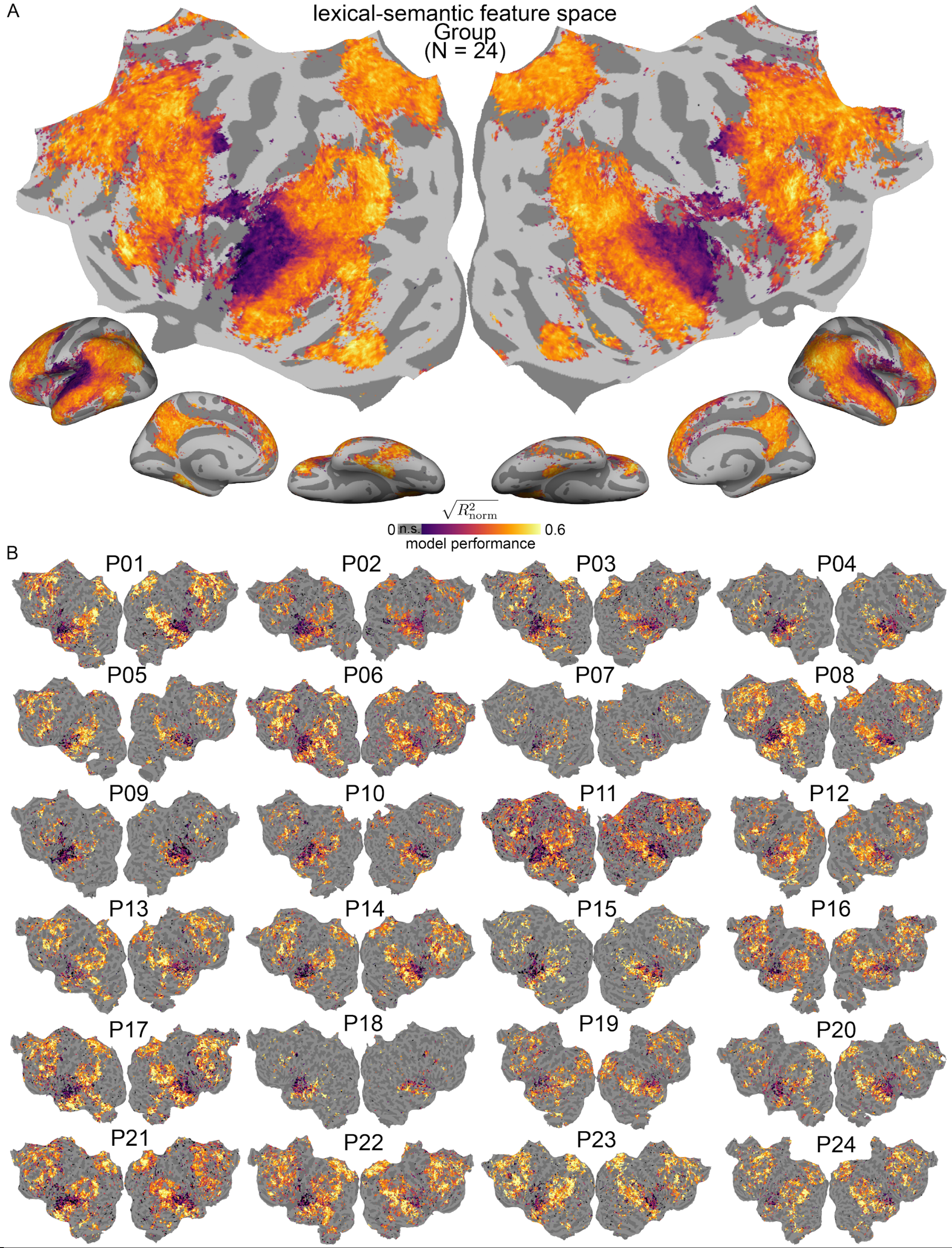


**Supplementary Figure 1. Maps of model performance for the lexical-semantic feature space.** To assess whether voxelwise encoding models can recover lexical-semantic information conveyed by narrative stories, model performance for the lexical-semantic feature space was statistically thresholded and displayed on the group-averaged and participant-specific maps. Participant-specific maps are thresholded at $p < 0.05$ (non-parametric permutation testing), and corrected for multiple comparisons with the Benjamini-Hochberg FDR procedure. [A] Group-averaged map showing only vertices in which at least six participants (25%) have significant model performance. [B] Participant-specific maps. These maps reveal that the lexical-semantic feature space significantly predicts brain activity in a distributed cortical network of temporal, parietal, and frontal areas. These results confirm that the voxelwise encoding models reliably capture representations of lexical-semantic information during narrative story-listening.

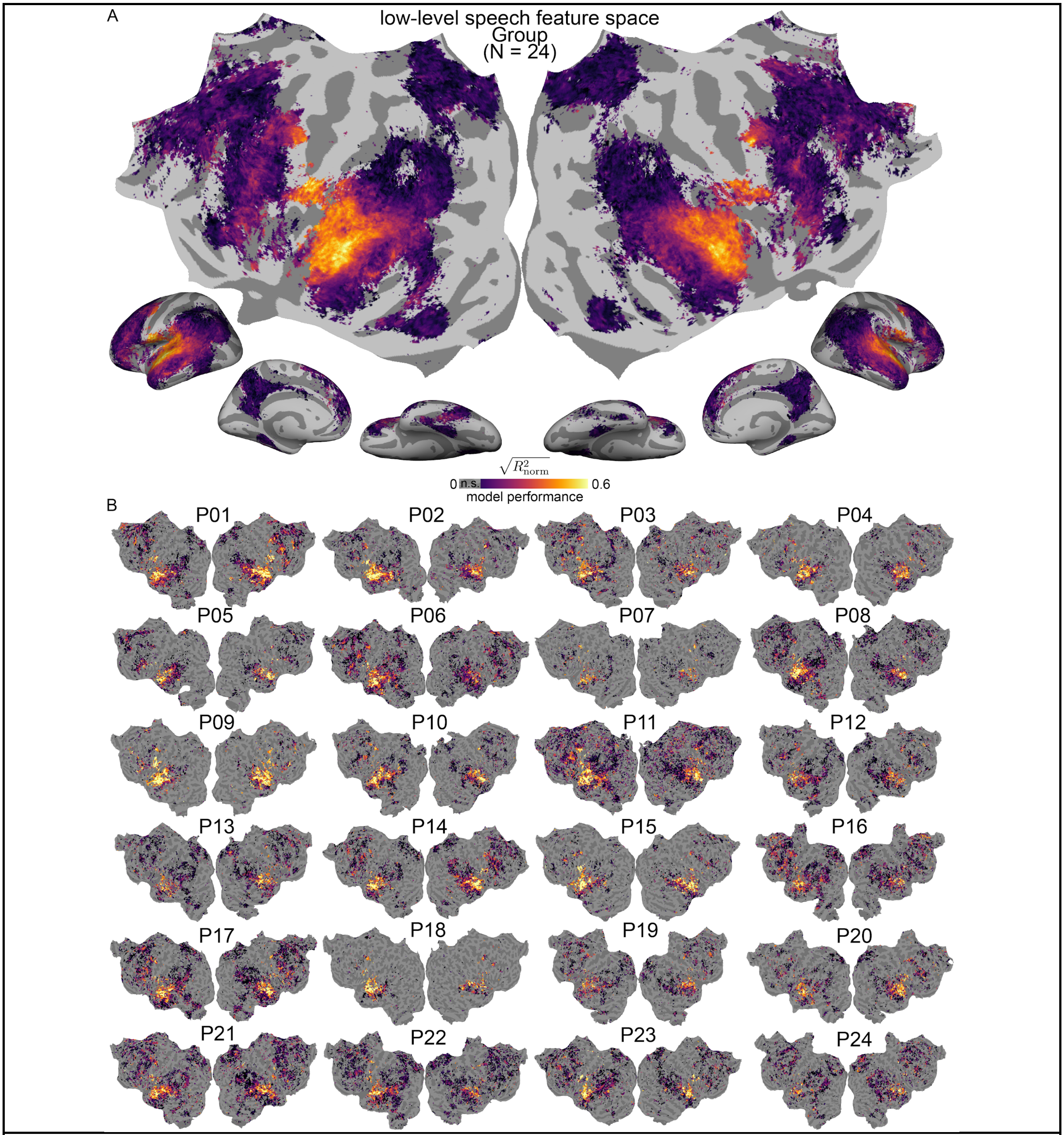


**Supplementary Figure 2. Maps of encoding model performance for the low-level speech feature space.** To account for variability in the speech pattern of different speakers, an additional low-level speech feature space was included in the banded ridge regression model (see Methods in Main Text). The maps in this figure show the prediction accuracy for the low-level speech feature space. [A] Group-averaged map. [B] Participant-specific maps. These maps reveal that the low-level speech feature space predicts brain activity in a narrow set of areas that include primary auditory cortex and speech-related areas. These areas overlap minimally with the areas predicted by the lexical-semantic feature space. The banded ridge approach used here can effectively disentangle representations of lexical-semantic information from representations of low-level speech information. This enables investigating lexical-semantic representations while taking into account variability in low-level speech information.

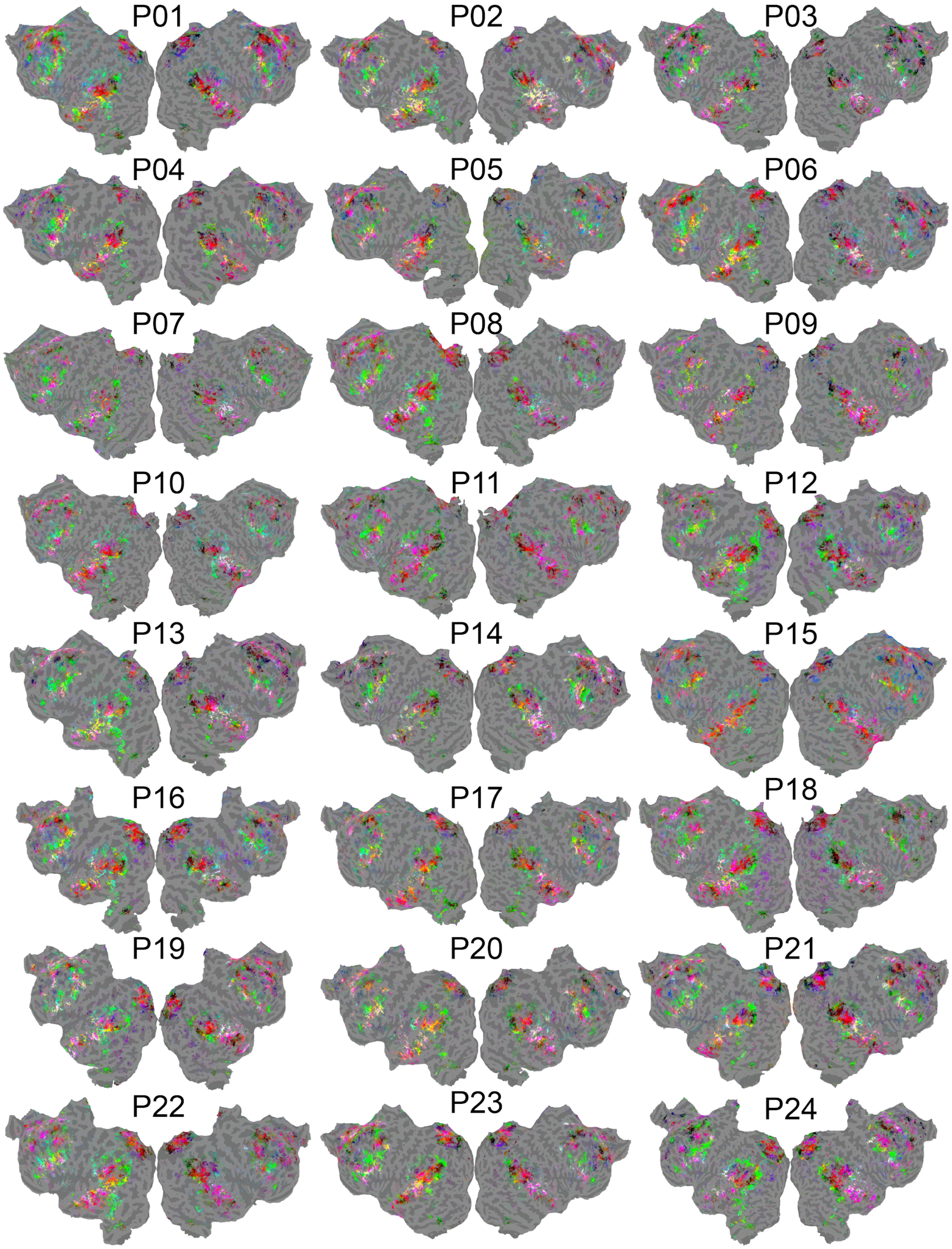


**Supplementary Figure 3. Participant-specific cortical maps of lexical-semantic representations.** To identify how lexical-semantic representations are spatially organized, and to determine whether they are variable across individuals, participant-specific model weights were examined on the cortical surface. Because the lexical-semantic model weights are 985-dimensional, it is difficult to visualize all dimensions on the cortical surface. Therefore, model weights were first projected onto a reduced three-dimensional lexical-semantic space (see Methods in Main Text). Then, the three dimensions were mapped onto the RGB color space to plot the selectivity of each voxel in this reduced lexical-semantic space. This visualization procedure produces a continuous colormap in which different colors indicate representations of different lexical-semantic concepts. Consistent with the results presented in the Main Text, the participant-specific maps in this figure highlight individual differences in lexical-semantic representations.

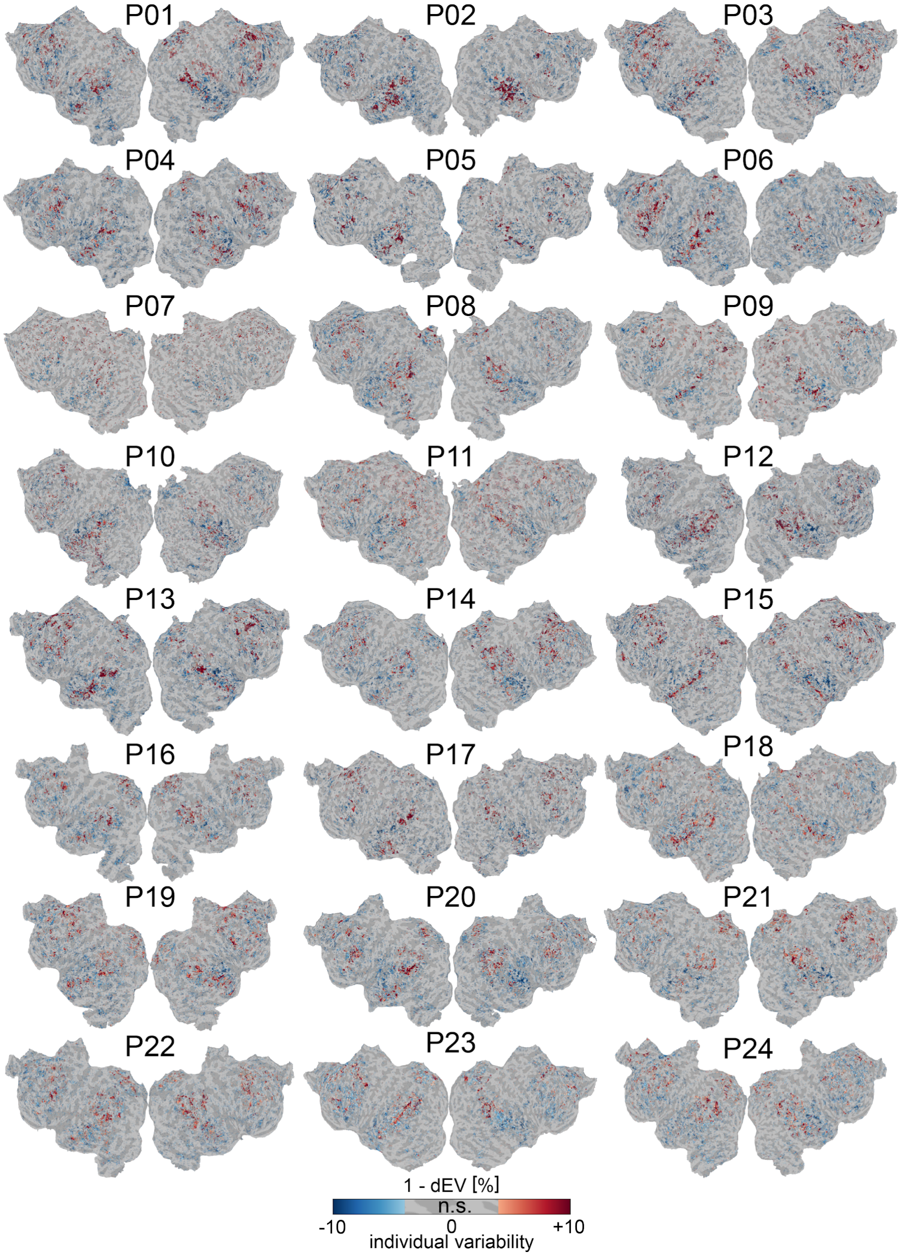


**Supplementary Figure 4. Spatial heterogeneity of individual differences in lexical-semantic representations.** Prior work suggests that individual differences in brain anatomy and function are spatially heterogeneous. However, whether this heterogeneity extends to complex lexical-semantic representations remains unknown. To address this question, we modified the dEV summary statistic to measure individual variability in each voxel. Because the dEV quantifies the variance explained by the group, individual variability is expressed as 1 - dEV. Individual variability maps were created separately for each participant. To retain only voxels with extreme individual variability values, maps were statistically thresholded at $p < 0.05$ using a jack-knife procedure (see Methods in Main Text). Blue indicates low individual variability; red indicates high individual variability. For each participant, values are expressed as relative change from the average. Consistent with the group-averaged map shown in Figure 3 in the Main Text, these participant-specific maps show that the largest individual differences are found in the TPJ, Precuneus, STS, and parts of the dorsomedial prefrontal cortex. These regions with high individual variability are thought to function as convergence zones, integrating modal and multimodal information to support conceptual knowledge. The smallest individual differences are found in the dorsal bank of the STS, in the ventral temporal lobe, and in portions of ventral Precuneus and prefrontal cortex. These regions with low individual variability are close to sensory areas such as primary auditory cortex, or they are predominantly tuned to concrete information such as visual-related concepts. These results demonstrate that associative and high-order areas exhibit the largest individual differences in lexical-semantic representations.

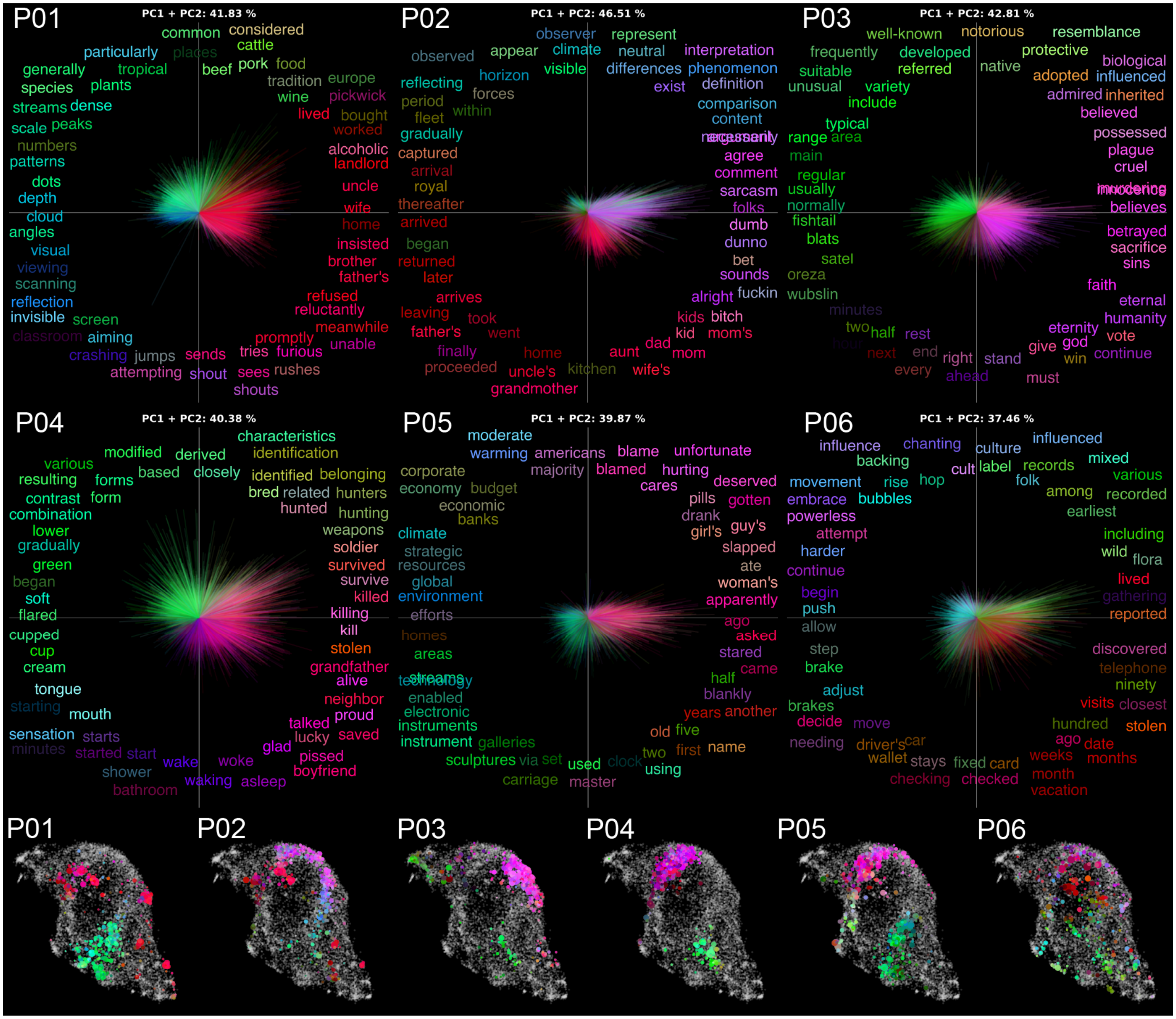


**Supplementary Figure 5. Individual differences in lexical-semantic representations reflect person-specific conceptual biases (P01-P06).** In the framework used in this work, individual differences are quantified using optimal transport to measure how each participant's lexical-semantic model weights differ from the group distribution of model weights. These differences reveal systematic biases towards specific lexical-semantic concepts, reflecting how each person's conceptual space differs from others. To identify the major dimensions underlying these person-specific conceptual biases, principal component analysis was applied separately to the weight differences of each participant. The first two principal components (PCs) are shown as orthogonal axes for participants P01-P06. Words from the *english1000* vocabulary that project strongly onto each dimension are displayed and colored according to the RGB color scheme from Figure 1 in the Main Text. Lines correspond to individual voxel biases towards specific lexical-semantic concepts. The line length is proportional to bias magnitude, and the color matches the closest word. The UMAP visualization at the bottom highlights the words in the *english1000* vocabulary (colored dots) with the largest projection onto the subspace spanned by the first two PCs. This visualization demonstrates that the subspaces of different participants span distinct portions of the lexical-semantic space, confirming that these individuals differ along fundamentally different conceptual axes. These results reveal that conceptual representations differ between individuals along lexical-semantic dimensions that are uniquely associated with each person. These person-specific dimensions reflect systematic biases in how different individuals represent conceptual information.

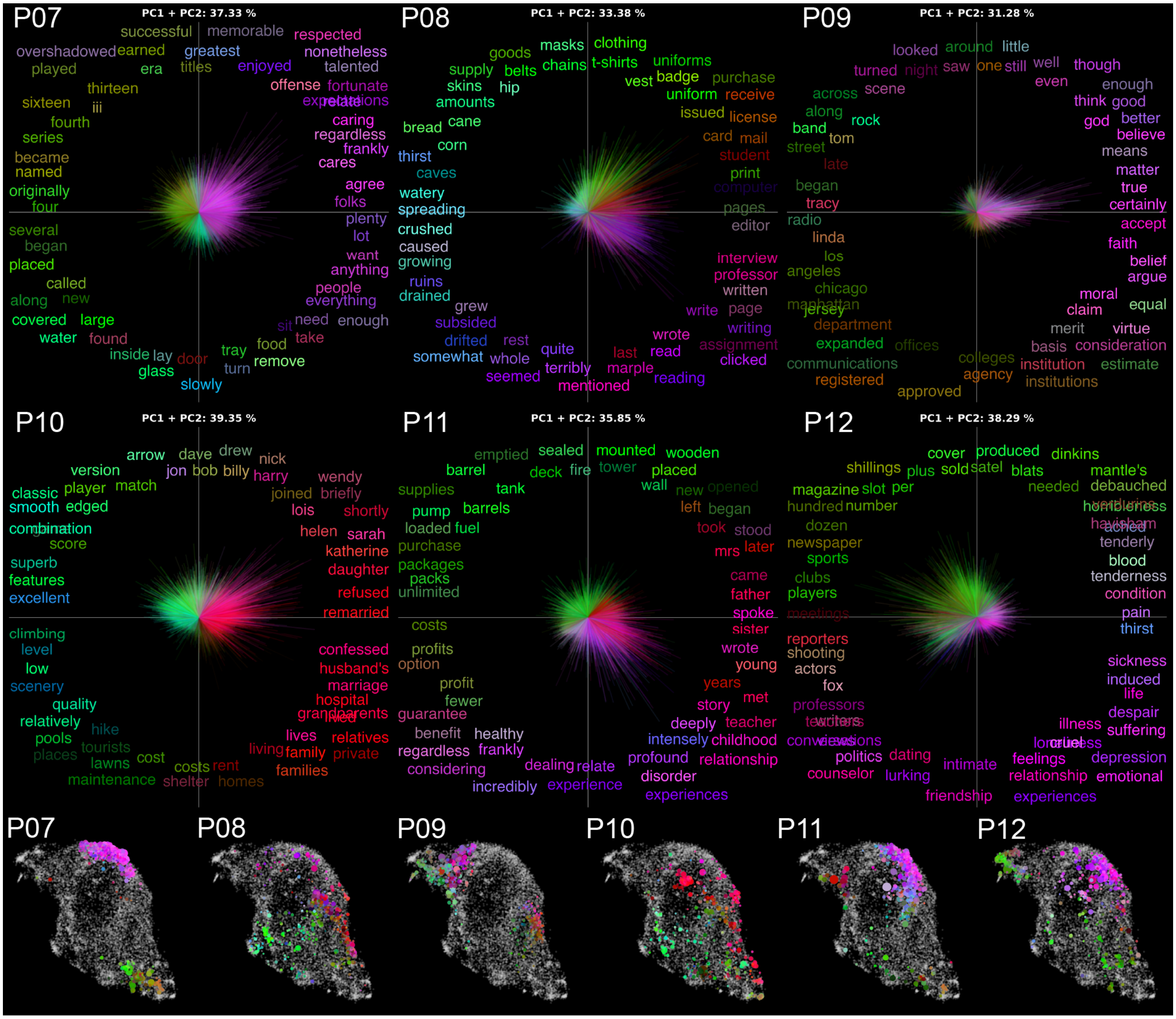


**Supplementary Figure 6. Individual differences in lexical-semantic representations reflect person-specific conceptual biases (P07-P12).** The first two principal components (PCs) underlying participant-specific conceptual biases and their UMAP visualizations are shown for participants P07-P12. See Supplementary Figure 5 for a detailed description of this figure.

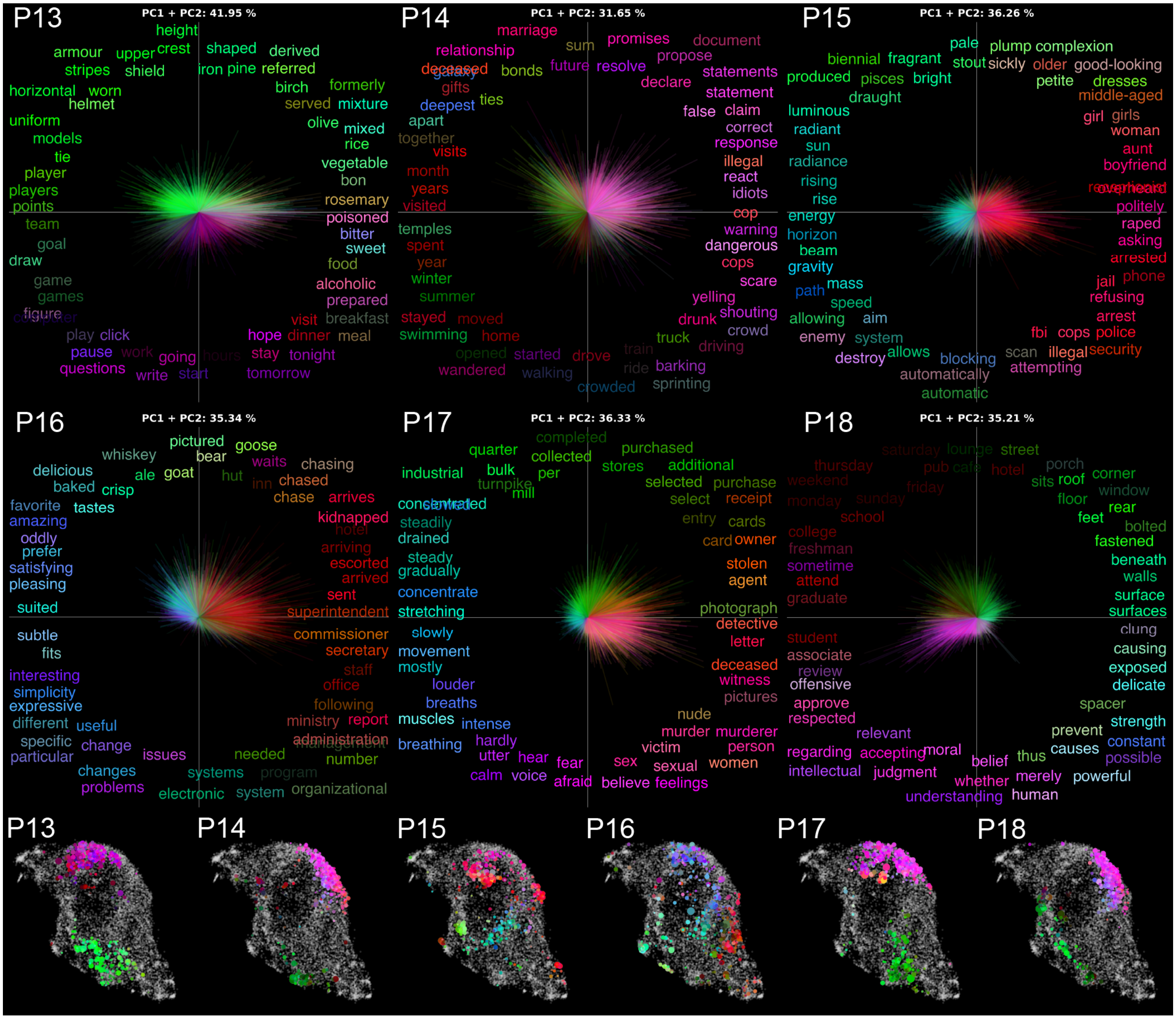


**Supplementary Figure 7. Individual differences in lexical-semantic representations reflect person-specific conceptual biases (P12-P18).** The first two principal components (PCs) underlying participant-specific conceptual biases and their UMAP visualizations are shown for participants P12-P18. See Supplementary Figure 5 for a detailed description of this figure.

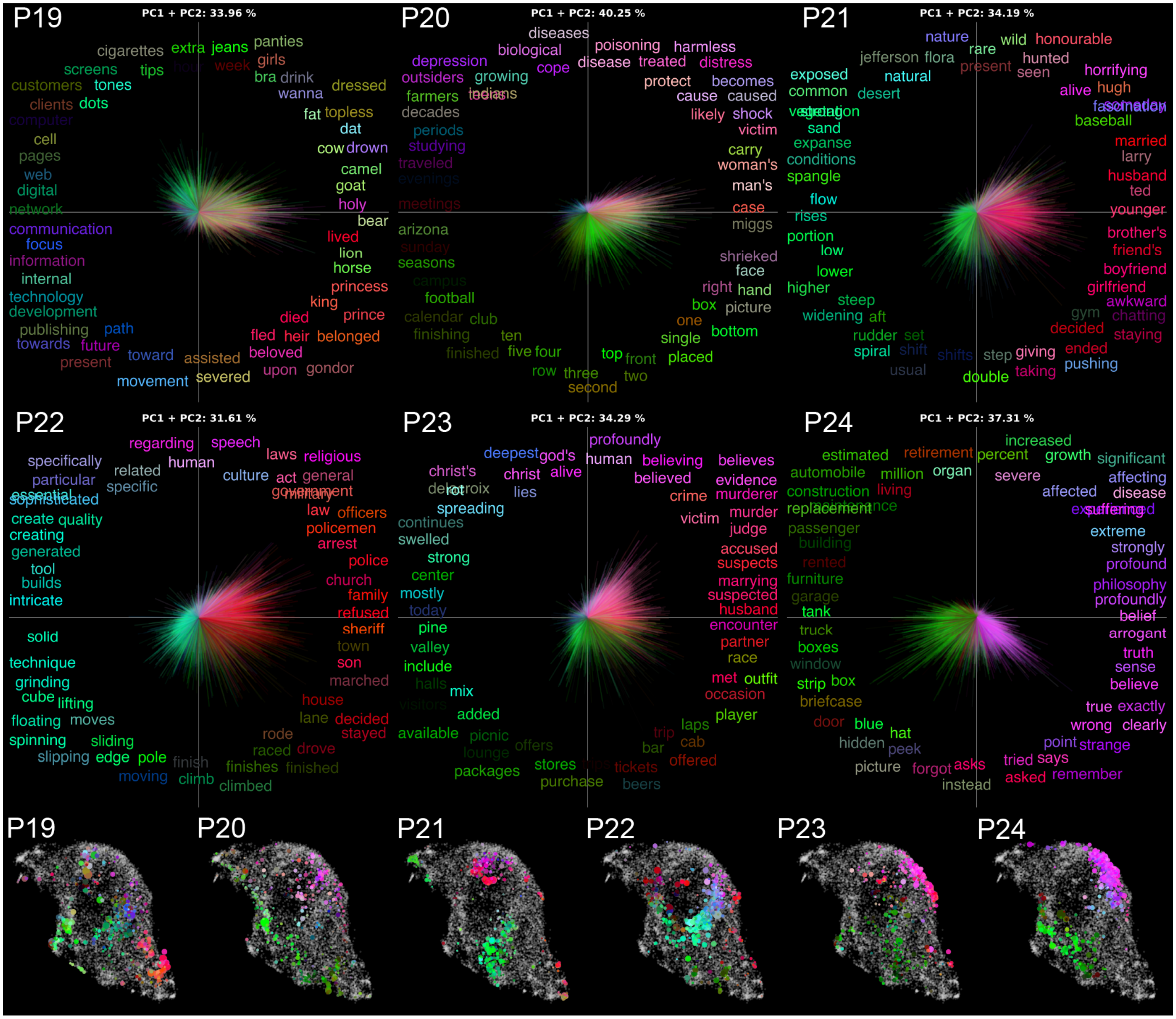


**Supplementary Figure 8. Individual differences in lexical-semantic representations reflect person-specific conceptual biases (P19-P24).** The first two principal components (PCs) underlying participant-specific conceptual biases and their UMAP visualization are shown for participants P19-P24. See Supplementary Figure 5 for a detailed description of this figure.

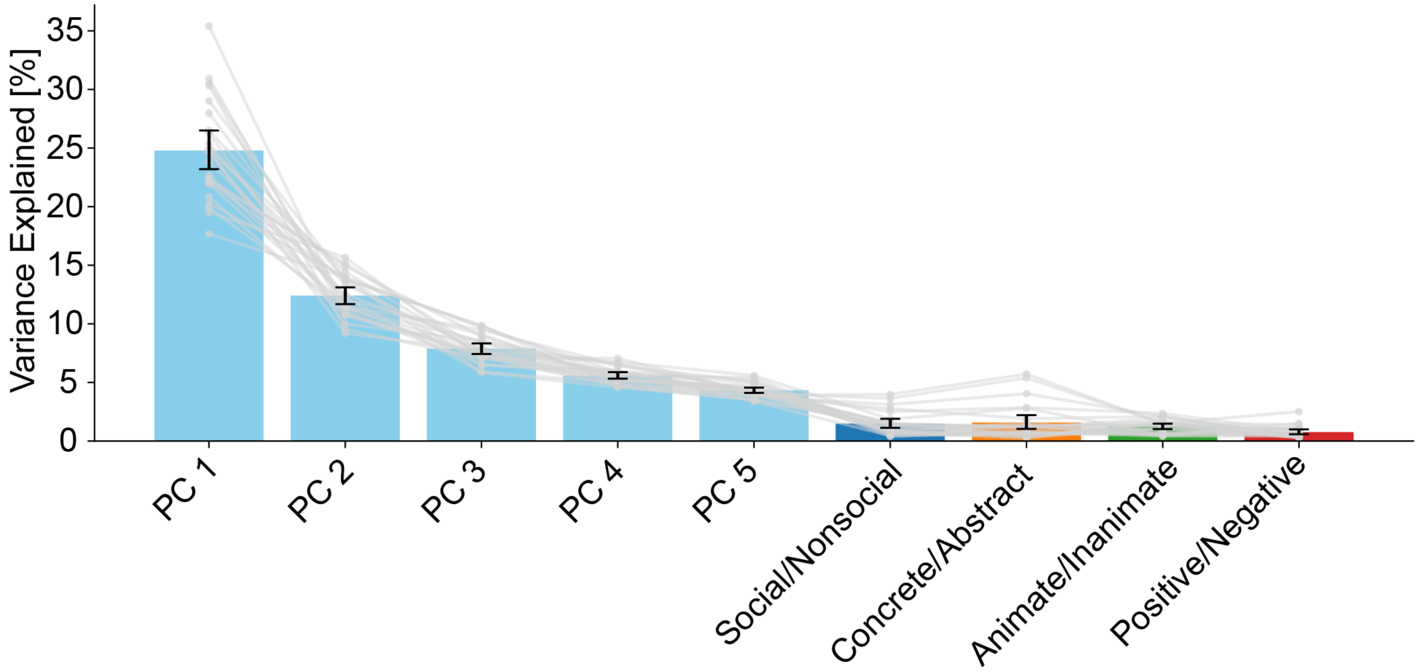


**Supplementary Figure 9. Canonical semantic axes do not explain participant-specific conceptual biases.** Prior research suggests that human conceptual knowledge is organized around several core semantic axes. To determine whether these semantic axes can explain the conceptual biases observed in this study, normed word lexicons were used to generate social/nonsocial, concrete/abstract, animate/inanimate, and positive/negative semantic axes in the *english1000* feature space (see Supplementary Methods). The fraction of variance explained by these axes was compared to the variance explained by the participant-specific lexical-semantic dimensions displayed in the Main Text and in Supplementary Figures 5-8. The bar plots indicate the average variance explained across 24 participants for the first five reliable PCs of each participant and for the four canonical semantic axes. Error bars indicate 95% bootstrapped confidence intervals. Gray lines indicate individual participants. These results show that the first two PCs of each participant explain on average more than 35% of variance in the conceptual biases. By contrast, the canonical semantic axes explain less than 2% of variance. Therefore, individual differences in lexical-semantic representations cannot be explained by canonical semantic axes such as social/nonsocial, concrete/abstract, animate/inanimate, or positive/negative valence. Instead, individual differences are best explained by dimensions that are specific to each participant, and reflect unique combinations of lexical-semantic concepts.

### Supplementary Tables

**Supplementary Table 1. Interpretation of the first lexical-semantic dimension underlying conceptual biases for each participant.** These labels were obtained with a data-driven procedure based on large language models (Summarize And Score, SASC; see Methods in Main Text for more details.)

| Participant | Negative | Positive |
|---|---|---|
| **P01** | observation, measurement, spatial, patterns | family, relationships, actions, places |
| **P02** | movement, time, events, royalty | informal communication |
| **P03** | faith, morality, humanity | variety, randomness |
| **P04** | descriptive, physical, attributes | life, emotions, crime |
| **P05** | environment, infrastructure, technology | actions, people, time, humor |
| **P06** | action, effort | discovery, rarity, history |
| **P07** | time, people, actions, numbers | communication, opinion, judgment |
| **P08** | environment, agriculture | communication, education, documentation |
| **P09** | music, people, places | belief, knowledge, reasoning |
| **P10** | outdoor adventure | family, relationships, events, crime |
| **P11** | economics, efficiency, quantity, cost | narrative, characters, actions, time |
| **P12** | health, suffering, treatment | media, sports, groups |
| **P13** | objects, sports, clothing, visuals | food, drink, preparation |
| **P14** | time, places, numbers | conflict, judgment, legality |
| **P15** | light, energy, movement, celestial | people, communication, relationships, politeness |
| **P16** | evaluation, preference | government, administration, time, locations |
| **P17** | movement, intensity | people, crime, communication |
| **P18** | structures, movement, surfaces | education, events, people |
| **P19** | communication, technology, organization | nobility, animals, life, death |
| **P20** | places, activities, time, groups | actions, possession, people, body |
| **P21** | geography, elevation, flow | people, relationships |
| **P22** | creation, process, quality | people, actions, institutions, conflict |
| **P23** | places, nature, structures | crime, relationships |
| **P24** | beliefs, understanding, emotions | objects, structures, transportation |

**Supplementary Table 2. Interpretation of the second lexical-semantic dimension underlying conceptual biases for each participant.** These labels were obtained with a data-driven procedure based on large language models (Summarize And Score, SASC; see Methods in Main Text for more details.)

| **Participant** | **Negative** | **Positive** |
|---|---|---|
| **P01** | actions, emotions, names | environment, geography, climate, agriculture |
| **P02** | science, observation, change | family, home, actions |
| **P03** | description, characteristics | time, actions |
| **P04** | everyday life, emotions | classification, characteristics |
| **P05** | emotions, health, substances | art, history, objects |
| **P06** | music, culture | travel, time, money |
| **P07** | objects, actions, conditions, locations | achievement, sports, success |
| **P08** | past events and emotions | clothing, accessories, equipment, goods |
| **P09** | organization, governance, education | everyday life |
| **P10** | living conditions, economy | names, sports |
| **P11** | objects, locations, actions, colors | mental processes, emotions |
| **P12** | human relationships | names, baseball, miscellaneous |
| **P13** | structure, materials, shape, position | communication, time, actions |
| **P14** | movement, places, time, nature | beliefs, commitments, assertions |
| **P15** | appearance, women | security, action, threat |
| **P16** | technology, communication, systems | actions, animals, food, settings |
| **P17** | human experience, emotions | business, location, quantity |
| **P18** | belief, reasoning, truth | places, movement, objects |
| **P19** | history, conflict | lifestyle, clothing, beverages |
| **P20** | health, risks | position, measurement |
| **P21** | life, death, nature, culture | movement, adjustment, quantity, objects |
| **P22** | society, perception, behavior | movement, position |
| **P23** | belief, morality, suffering | leisure, commerce |
| **P24** | health, impact, time, economy | actions, objects, appearances |

**Supplementary Table 3. Words selected according to normed lexicons to generate four core semantic axes.**

| Category | Words |
|---|---|
| **Social** | friendship, relationship, people, romance, marriage, political, family, boyfriend, friend, mother, sister, festival, meeting, mommy, society, conversation, politics, politician, civilization, funeral, mankind, friendly, argument, communicate, rumor, community, citizen, discussion, language, cult, culture, sexual, thank, loyal, colleague, sibling, volunteer, kinship, husband |
| **Nonsocial** | horizontal, sleeve, button, birch, forearm, windshield, flannel, debris, pebble, silvery, broomstick, metal, particle, bracelet, axe, tiger, handkerchief, syllable, hillside, lantern, plaster, pantry, blank, ceiling, bulk, cupboard, needle, flask, cotton, whole, rack, stair, colon, bowel, initial, can, curtain, barren, bonnet |
| **Abstract** | although, whatsoever, belief, perhaps, though, terribly, inexplicable, infinitely, wherever, hope, ultimately, nonetheless, meanwhile, especially, somewhat, somehow, basically, incomprehensible, as, enough, despite, possibility, luck, disbelieving, forever, infinity, necessarily, indefinite, thus, oblivion, bliss, actually, sincerely, suppose, ideal, theoretical, evidently, seldom, anything |
| **Concrete** | shawl, horse, clock, bird, knee, eyelid, gravel, granite, ball, frog, camera, apple, ear, comb, binoculars, flashlight, whisky, bed, passport, bat, carrot, bean, water, pig, sand, goat, leaf, turtle, eagle, lemon, ladder, mattress, penis, plank, snake, magazine, bike, spear, horn |
| **Animate** | officer, son, frog, aunt, cousin, elephant, cat, baby, animal, waiter, waitress, bride, citizen, farmer, brother, kid, pony, captain, roommate, pope, goose, singer, dog, mother, father, actress, hostage, physician, monkey, tortoise, girl, wife, bee, daughter, hostess, lion, customer, cow, boyfriend |
| **Inanimate** | whip, lamp, fork, handkerchief, platform, instrument, salt, bench, costume, ribbon, van, ladder, stool, envelope, flag, bra, kettle, bucket, shovel, wallet, mask, pole, boot, train, coin, stove, drum, shirt, jacket, quarter, napkin, bottle, glass, mug, jug, string, needle, rifle, compass |
| **Positive** | clap, providing, share, enjoy, comfort, venerable, successful, intimate, brilliant, eagerness, dove, bless, lover, proud, bloom, glow, engaged, happy, salary, perfect, treasure, star, grow, healing, friendly, pay, infinity, champion, victory, faith, passionate, familiarity, cradle, readiness, radiance, grant, respect, marriage, glory |
| **Negative** | lose, terrorist, horror, murderous, howl, disaster, thief, wreck, catastrophe, murder, attacking, slaughter, bang, distress, traitor, jealousy, poverty, bitch, abuse, cruelty, desert, horrid, ruined, powerless, curse, disease, poisoned, theft, mad, unlucky, painful, illegal, hell, confined, terrible, demon, nasty, scar, robbery |
